# Fundamental limits on patterning by Turing-like reaction-diffusion mechanisms

**DOI:** 10.1101/2025.05.27.656324

**Authors:** Daniel Muzatko, Bijoy Daga, Tom W. Hiscock

## Abstract

Turing’s longstanding reaction-diffusion hypothesis explains how molecular patterns can self-organize *de novo* in otherwise homogeneous tissues. However, whilst simple Turing models can qualitatively recapitulate patterning *in silico*, they are often highly simplified approximations of the molecular complexity operating *in vivo*. Here, we investigate significantly more complex reaction-diffusion models that seek to directly capture the mechanisms involved in intercellular signalling. Rather than resulting in minor, quantitative differences, we find that these more realistic systems display qualitatively distinct behaviours compared to simple Turing models. By combining large-scale simulations with formal mathematical proofs, we show, rather generally, that patterning is strongly constrained by the extracellular interactions in the system but is relatively insensitive to the intracellular dynamics assumed. When applied to the activator-inhibitor paradigm, we find a broader repertoire of self-organizing circuits than previously recognized, including some which are unexpectedly robust to parameters. Beyond these examples, we have packaged our highly performant numerical methods into a freely available and easy-to-use software pipeline, ReactionDiffusion.jl, that allows arbitrarily complex reaction-diffusion systems to be simulated at scale.

## INTRODUCTION

Symmetry breaking is one of the most fundamental processes in developmental biology, allowing otherwise homogeneous tissues/embryos to self-organize into complex patterns and structures. The ability of early embryos to form patterns *de novo* has long been recognized, dating back to some of the earliest experiments in embryology in the late 1800s (Driesch, 1892; Roux, 1888). Beyond early embryos, self-organization also occurs in many developing tissues, as evidenced by the ability of diverse organoid systems to self-organize patterns *in vitro* (Eiraku et al., 2011; Ishihara and Tanaka, 2018; Meinhardt et al., 2014; Merle et al., 2024; Simunovic et al., 2019). Some of the most well-studied paradigms of self-organization involve periodic (i.e., repeating) patterns (Hiscock and Megason, 2015; Kondo and Miura, 2010; Marcon and Sharpe, 2012; Raspopovic et al., 2014; Sick et al., 2006), in which initially uniform tissues spontaneously break symmetry to generate identical/similar structures at regular spatial intervals (e.g., teeth, hair, leaves).

Decades of theoretical work – initiated by Alan Turing in the 1950s (Turing, 1952) – has advanced the hypothesis that these self-organizing patterns are the result of reacting and diffusing chemical signals. *In vivo*, these reaction-diffusion systems can be implemented by intercellular feedback circuits of secreted signalling molecules that diffuse in the extracellular space. In the canonical Turing model (as well as subsequent Gierer-Meinhardt models (Gierer and Meinhardt, 1972; Meinhardt and Gierer, 1974)) there are two such diffusible molecules – an activator and its (more rapidly diffusing) inhibitor (Figure 1A). These activator-inhibitor systems can self-organize *in silico* from a homogeneous initial state into a diversity of eventual patterns, often periodic patterns such as dots or stripes. Moreover, putative activator/inhibitor pairs have been proposed for a variety of tissues and signalling pathways, and *in silico* Turing models can be made to qualitatively recapitulate patterns formed *in vivo* (Grall et al., 2024; Kondo and Asai, 1995; Raspopovic et al., 2014; Sick et al., 2006).

**Figure 1.**
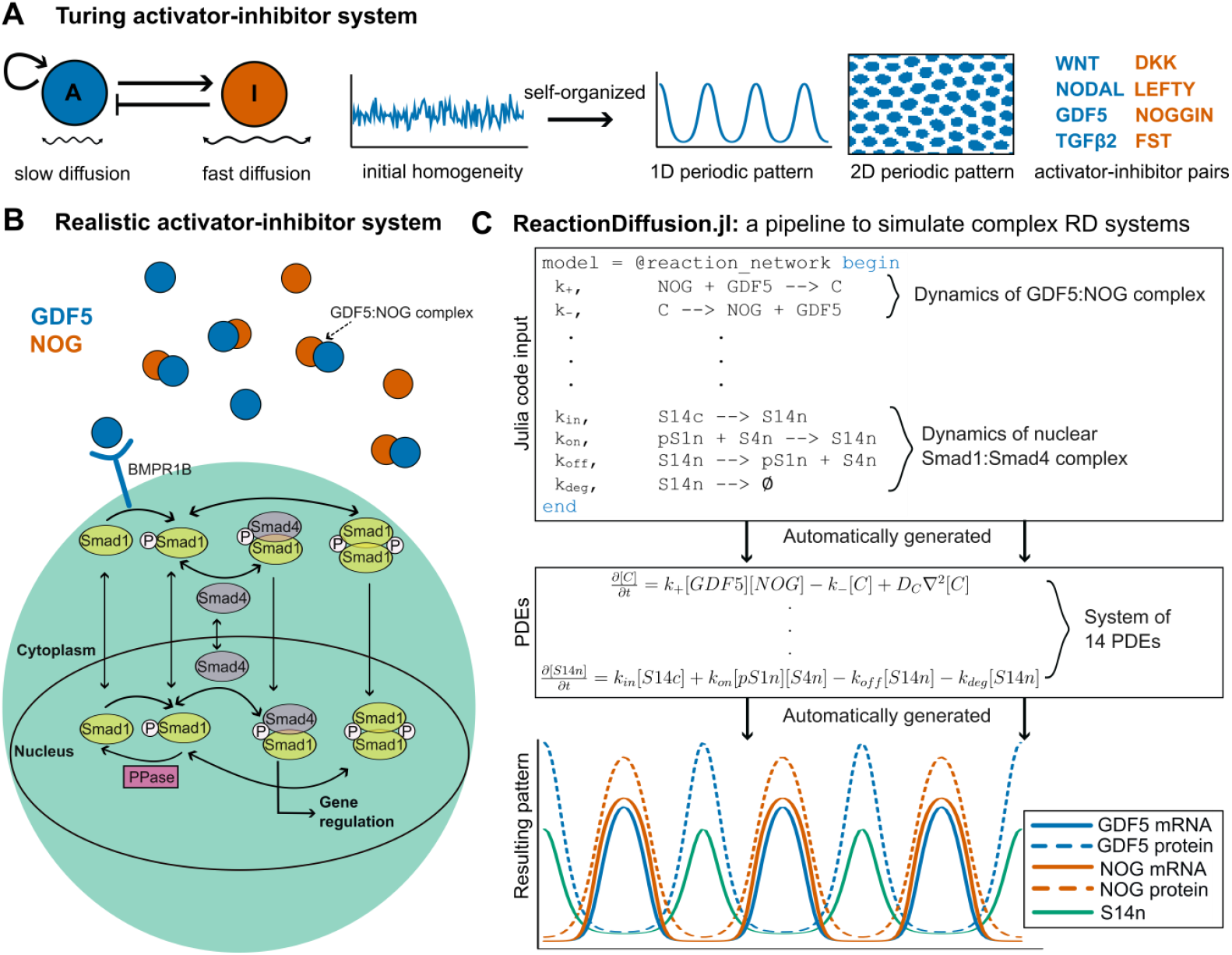
Self-organization in activator-inhibitor systems. (**A**) Simplified reaction-diffusion models that comprise two diffusible molecules – an activator and its own inhibitor – can self-organize repeated patterns *in silico* (Gierer and Meinhardt, 1972; Turing, 1952). (**B**) Schematic of complex intercellular interactions involved in an example activator-inhibitor system *in vivo* (Grall et al., 2024; Schmierer et al., 2008). **(C**) Outline of ReactionDiffusion.jl, a computational pipeline to efficiently simulate complex reaction-diffusion systems. The model shown corresponds to the interactions schematized in (**B**). Model assumptions are written in an intuitive syntax that reflects biochemical interactions. Models are automatically converted to partial differential equations, and then automatically simulated until reaching a stationary pattern.

However, Turing models are often highly simplified, approximating complex mechanisms by a handful of equations that loosely mirror the regulatory interactions present *in vivo*, and neglect the essential roles that cellular and/or mechanical interactions may be playing (Hiscock and Megason, 2015; Ho et al., 2019; Nakamasu et al., 2009; Shyer et al., 2017). Moreover, when modelling pure reaction-diffusion systems *in silico*, they are highly sensitive to parameters and often only form patterns in a narrow and biophysically unrealistic region of parameter space (Maini et al., 2012; Scholes et al., 2019). It is thus unclear whether a molecular mechanism, alone, could ever be capable of robust patterning, casting doubt on the relevance of Turing’s hypothesis *in vivo*.

Despite this, recent experimental data are beginning to pinpoint specific patterns that are generated by definitively molecular (i.e., reaction-diffusion) mechanisms. The clearest examples draw from tissues that undergo minimal morphogenesis during initial patterning, such as the developing hair follicles (Sick et al., 2006) and fingerprints in the mammalian epidermis (Glover et al., 2023), or the patterning of joints within the digits (Grall et al., 2024). In these cases, a self-organized molecular prepattern is the first sign of symmetry breaking and precedes any overt changes in cell density or tissue morphology, strongly implicating a reaction-diffusion mechanism.

At the same time, theoreticians have started to consider reaction-diffusion systems that are considerably more complex than simple activator-inhibitor schemes e.g., by incorporating more than two diffusible molecules. High throughput, systematic analyses of 3- and 4-node networks have revealed a wide array of regulatory architectures capable of periodic patterning beyond simple 2-component Turing models (Diego et al., 2018; Haas and Goldstein, 2021; Marcon et al., 2016; Scholes et al., 2019; Shaberi et al., 2025; Zheng et al., 2016), suggesting that patterning may involve more than just one activator and one inhibitor *in vivo*. However, whilst containing more diffusible elements, the models are still simplified network-level abstractions which approximate molecular interactions by simple mathematical expressions (e.g., linear or hill functions), rather than directly capturing the nature of developmental signalling *in vivo*. Moreover, many of the identified circuits remain highly fragile to parameters *in silico* (Scholes et al., 2019).

Here, instead of adding more activators/inhibitors, we seek to build models of classic Turing mechanisms (i.e., 1-activator, 1-inhibitor) that more realistically capture the mechanisms of intercellular signalling. We move beyond idealized 2-component representations of these systems, and instead explicitly consider the multi-step nature of pathway activation (i.e., signal transduction), transcriptional regulation (i.e., feedbacks) and the diverse biochemical mechanisms of pathway inhibition (e.g., some inhibitors sequester ligands, whereas others compete for receptor binding). To deal with the substantial increase in model complexity that this entails, we developed a highly performant and freely available computational pipeline (ReactionDiffusion.jl) that simulates complex reaction-diffusion systems at scale and at speed. By combining large scale numerical simulations with formal mathematical proofs, we uncover fundamental limits on the ability of such activator-inhibitor circuits to break symmetry, which apply rather generally across many types of signalling pathways and intracellular dynamics.

Rather than all behaving similarly – as implicitly assumed in simple 2-component Turing models – we have identified eight distinct classes of activator-inhibitor system, defined by the type of inhibitory mechanism and regulatory feedbacks present. This suggests that self-organized patterning is possible with a more flexible regulatory logic than previously thought and reveals several pattern-forming motifs not predicted by simpler models. Moreover, we predict that patterning is achievable with biologically feasible parameter values, and in some cases, can be made robust to variations in any single biophysical/biochemical parameter. Thus, by adding complexity and realism to our models, we have revealed a more widespread ability of reaction-diffusion systems – even those that comprise only a single activator and single inhibitor – to self-organize patterns *de novo*.

## RESULTS

### ReactionDiffusion.jl: a pipeline to efficiently simulate complex reaction-diffusion systems

Intercellular feedbacks in reaction-diffusion systems couple cell states across space and time within developing tissues. These interactions can be coarse grained via a continuum approximation into systems of partial differential equations (PDEs) that describe the spatiotemporal dynamics of signalling. Whilst simple to write down, the complexity of our models (large systems of numerically stiff PDEs) makes them highly challenging to simulate. We therefore developed a suite of new computational tools to simulate these complex reaction-diffusion systems at scale.

We have built an end-to-end pipeline in the performant *SciML* ecosystem that leverages an expressive domain-specific-language, state-of-the-art numerical solvers, and automated multicore parallelization (Loman et al., 2023; Ma et al., 2021; Rackauckas and Nie, 2019, 2017). Models are first specified in an intuitive syntax that naturally reflects the regulatory interactions involved in signalling (Figure 1B, C). Mass action kinetics are automatically prescribed for the specified biochemical reactions, and user-defined functions can be used to parameterize other reaction rates (e.g., transcriptional feedbacks). Once specified, the model is then *automatically* converted to a system of corresponding PDEs, and then *automatically* simulated to predict the resulting dynamics, all within a single line of code (Figure 1C). We have implemented highly-performant, adaptive time-stepping solvers that allow systems of 10+ PDEs to be simulated interactively on a standard desktop machine (see Supplementary Materials).

We have also implemented Turing instability analysis into our pipeline, mirroring the approaches used in (Diego et al., 2018; Scholes et al., 2019) to predict whether a reaction-diffusion system can break symmetry (Figure S1A). Again, this is fully automated, going from an intuitive model description to a high-throughput parameter screen in just a single line of code (Figure S1B). We have demonstrated the efficacy of our approach by correctly identifying pattern-forming parameter sets for 4 separate Turing systems (2-4 components) with known ground truths (Figure S1C-F). By employing multithreaded parallelization and symbolic computing methods, we can accelerate these computations and are able to screen millions of parameter sets per minute on a standard desktop machine.

Together, these form a fully end-to-end pipeline for efficiently simulating and screening complex reaction-diffusion systems. These tools can be applied to any type/size of reaction-diffusion system and are fully automated, meaning that a detailed knowledge of numerical solvers or fine tuning of algorithm parameters is not required. We have made our pipeline freely-available as an open-source Julia package with extensive documentation and tutorials (https://github.com/hiscocklab/ReactionDiffusion.jl).

### Deriving general limits on pattern formation by reaction-diffusion systems

Armed with our new pipeline, we are now able to simulate reaction-diffusion models with sufficient complexity to more accurately mirror intercellular signalling mechanisms. However, despite the significant acceleration in simulation time, the curse of dimensionality still constrains our ability to screen large systems of PDEs. For example, the GDF5 signalling model in Figure 1 has 20 free parameters and 14 components. Whilst we can simulate this rapidly for a single parameter set, to exhaustively screen through all combinations remains computationally unfeasible. Therefore, we sought strategies to approximate our detection of Turing instabilities to render comprehensive screens tractable, allowing us to derive generally applicable limits regardless of parameter values (many of which are often not quantified *in vivo*).

For a Turing-like instability to arise, there must be some small perturbations with wavenumber *q* (equivalently, periodic patterns with wavelength *λ* = 2*π*/*q*) which destabilize an initially homogeneous state (J.D. Murray, 2008). One simplification that has been used previously is to examine the determinant of the linearised reaction-diffusion matrix, **M**(*q*^2^) ≡ **J** − *q*^2^**D** (Diego et al., 2018; Smith and Dalchau, 2018), where **J** is the Jacobian of the reaction terms and **D** is a diagonal matrix corresponding to diffusion. The Turing instability criterion requires det (**M**(*q*^2^)) to change sign twice in the range *q* ∈ [0, ∞) (Smith and Dalchau, 2018), which we hereafter refer to as the necessary condition **𝒩** (Figure 2A).

**Figure 2.**
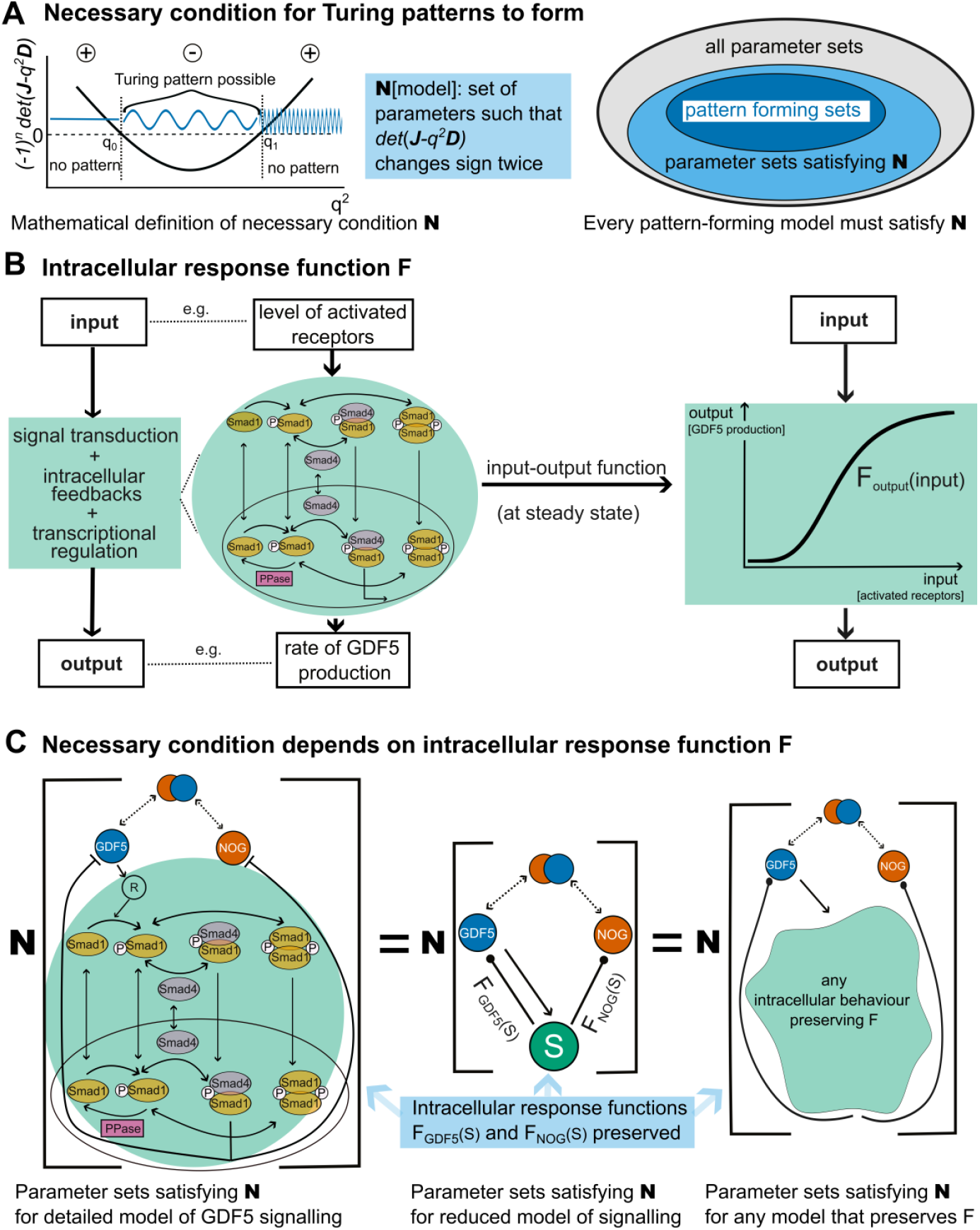
Necessary conditions for any reaction-diffusion system to break symmetry. **(A)** Schematic of a necessary condition for patterning, **𝒩**, which requires the determinant of the linearized reaction-diffusion matrix to change sign for some finite range of pattern wavevectors, *q*. (**B**) The intracellular response function, F_output_(input), approximates intracellular interactions (e.g., signal transduction, transcriptional feedbacks) by a single input-output function per pathway, assuming that all intracellular components have reached steady state. (**C**) The necessary condition, **𝒩**, of a reaction-diffusion system can be computed without full knowledge of its intracellular dynamics, provided the steady-state intracellular response function F_output_(input) is known.

When considering detailed reaction-diffusion models (e.g., like Figure 1B), we noticed that their structure greatly simplifies the calculation of **𝒩** (see Supplementary Materials). In general, signalling pathways comprise elements which diffuse between cells (e.g., extracellular ligands and inhibitors; we label these as *D*); elements which directly interact/bind to the diffusible elements (e.g., receptors; we label these as *R*); and elements which are confined within the cell itself (e.g., intracellular signal transduction elements; we label these as *I*). We may split **M**(*q*^2^) accordingly into blocks, so for example **M**_***DD***_(*q*^2^) ≡ **J**_***DD***_ − *q*^2^**D**_***DD***_ represents interactions/diffusion of the extracellular components *D*, whereas **M**_***IR***_(*q*^2^) ≡ **J**_***IR***_ reflects how the intracellular elements *I* are regulated by (activated) receptors *R*. Whilst the intracellular components, *I*, add significant complexity to the models, they do not contribute to intercellular diffusion (i.e., **D**_*II*_ = **0**) and are not *directly* affected by the extracellular species (i.e., **J**_*ID*_ = **0**). Using Schur’s lemma, we therefore find that the determinant of **M** may be expressed in a simpler form:

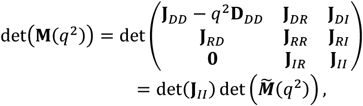

where 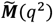 (*q*^2^) is defined as:

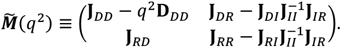

Since det(**J**_*II*_) is a constant, it follows that the necessary condition, **𝒩**, may be computed by considering a much simpler system, 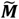, which has precisely the same extracellular interactions (**J**_*DD*_, **J**_*RD*_) and diffusion (**D**_*DD*_), but with all receptors and intracellular components removed and replaced with a single, non-diffusing component per signalling pathway (we label these *effective* intracellular components as *S*). The overall effect of intracellular signalling on the diffusible elements is then captured by 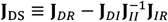, and the dynamics of intracellular signalling is represented by 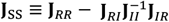. At first glance, the computation of these effective interactions remains challenging and appears to depend on the full complexity of the intracellular components. However, we may use Lemma 2.1 from (Smith and Dalchau, 2018), to show that these terms can be computed by simply assuming that all intracellular components, *I*, reach quasi-steady state i.e., do not change over time. Therefore, we may rigorously approximate the intracellular dynamics as a “black box” that takes in inputs (e.g., levels of activated signalling receptors) and translates these into outputs (e.g., production rates of signalling ligands and inhibitors), with each steady-state input-output relation (Figure 2B) quantified by a corresponding intracellular response function, F_output_(input). The advantage of this formulation is that the necessary condition **𝒩** may now be computed by considering the intracellular response functions alone without a detailed characterization of the intracellular dynamics. Moreover, we show that **𝒩** is also sufficient to predict the possible wavelengths of pattern that can form, as well as the phase relations between gene expression domains i.e., whether they are in- or out-of-phase (see Supplementary Materials).

There are multiple ways to measure F_output_(input) in experiments. The most direct would be to experimentally vary the input *in vitro* (e.g., via dosed application of exogeneous ligand to low density cultures) and then quantify the resulting output (e.g., via qPCR quantification of relevant transcript levels). Alternatively, one could use *in vivo* correlations to quantify how natural variations in inputs (e.g., levels of pathway activation measured via immunofluorescence) lead to changes in output levels (e.g., gene expression levels measured via fluorescent in situ hybridization). Moreover, even if F_output_(input) cannot be measured directly, it’s salient features can often be estimated or at least bounded. We later show that the ability of reaction-diffusion systems to break symmetry depends only on the normalized sensitivity of the intracellular response functions (Paulsson, 2004); this is defined as the % change in output for a given % change in input, i.e., H_input→output_ ≡ (ΔF_output_/F_output_)/(Δinput/input).

These normalized sensitivities, H_input→output_, have an intuitive interpretation – they can be viewed as the effective hill coefficients of the response functions (Figure 3A). These can either be directly estimated from data e.g., (Park et al., 2019), or bounded by biophysical arguments, e.g., (Martinez-Corral et al., 2024).

**Figure 3.**
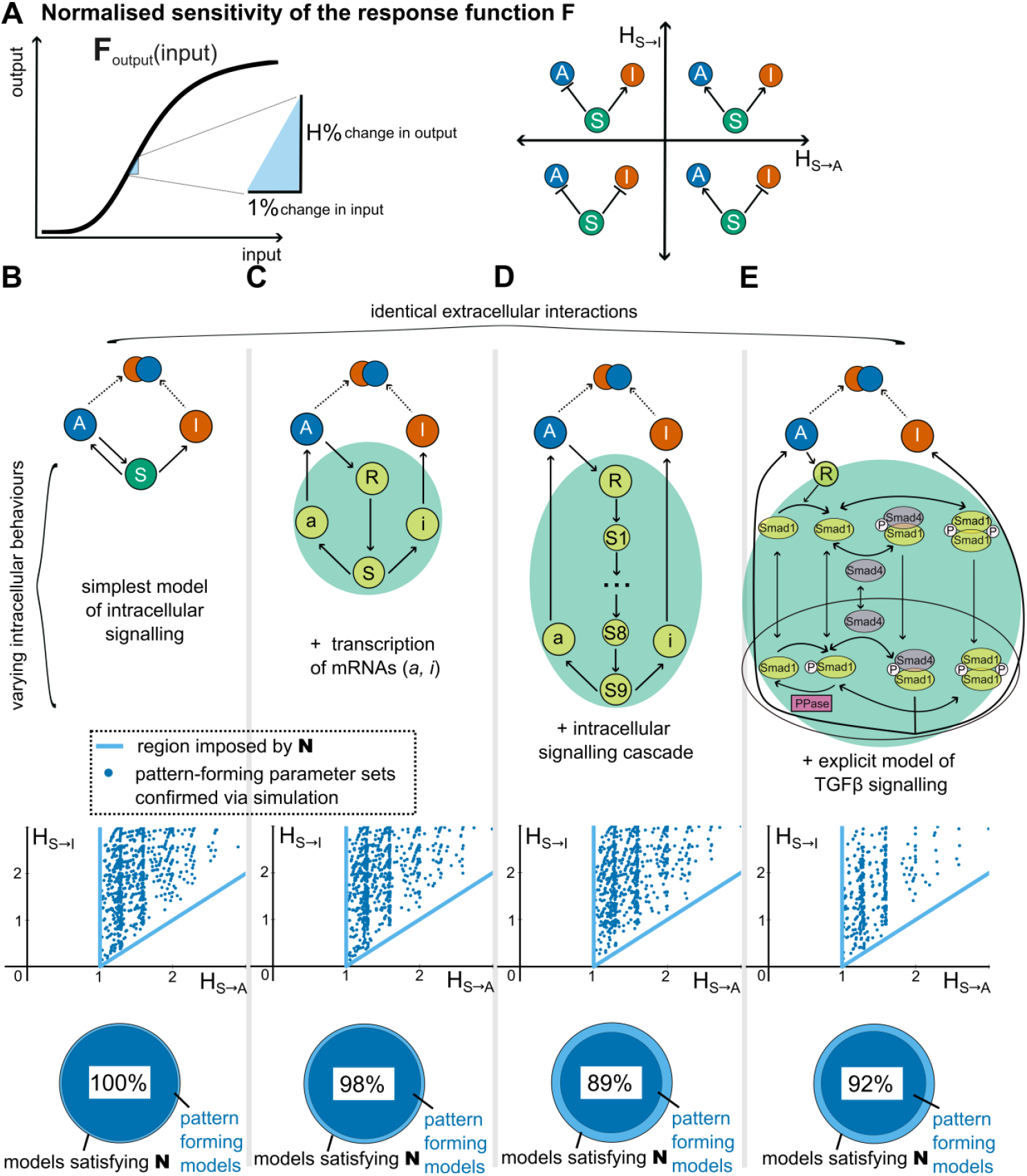
The ability of activator-inhibitor systems to form patterns is relatively insensitive to their intracellular dynamics. (**A**) The normalized sensitivity, H_input→output_, approximates the effect of a given intracellular feedback by a single number, which reflects how the output responds to changes in the input. H > 0 corresponds to activating, and H < 0 to repressive interactions. (**B**-**E**) Analysis of activator-inhibitor systems with the same extracellular but different intracellular interactions. All models assume that the extracellular inhibitor directly binds to the signalling ligand targeting it for degradation. A variety of intracellular behaviours are considered (schematics, upper row). Regardless of the intracellular network, we find that the sensitivity parameters are subject to the same constraints across all models (light blue lines, middle row) which is confirmed via numerical simulation (blue dots, middle row). By simulating many combinations of parameters for each model, we find that the set of pattern-forming parameters can be reasonably approximated by the necessary condition **𝒩** alone (~90% agreement, lower row), and is thus similar across all models despite differences in their intracellular regulation.

Taken together, these results provide strict limits on pattern formation (via the necessary condition **𝒩**) that can be derived without quantifying the full dynamics of intracellular signalling and regulation. Thus, we can now derive general limits on patterning that will be broadly applicable regardless of the precise intracellular feedback mechanisms – or intracellular parameters – at play (Figure 2C).

### The ability of reaction-diffusion systems to form patterns is relatively insensitive to their intracellular dynamics

Whilst the condition **𝒩** places strict constraints on pattern-forming parameters, it is not clear how tight – and thus how informative – these constraints are. Equivalently, **𝒩** defines a necessary condition for patterning, but it is unclear how close this is to a necessary *and* a sufficient condition. The complexity of intracellular dynamics we consider makes an analytical derivation of the sufficient conditions intractable. Therefore, we instead investigate this question numerically using our ReactionDiffusion.jl pipeline to determine pattern-forming parameters (i.e., those that satisfy both necessary and sufficient conditions). To begin with, we consider perhaps the simplest mode of pathway inhibition, in which the inhibitor directly binds to the signalling ligand thus preventing it from activating receptors and eventually targeting it for degradation. Keeping this extracellular component identical, we then consider a variety of intracellular dynamics: a single, effective intracellular signal (Figure 3B); incorporation of mRNA transcription (Figure 3C); a multi-step intracellular signalling cascade (Figure 3D); and an explicit model of GDF5 signal transduction (Nunns and Goentoro, 2018; Schmierer et al., 2008) (Figure 3E). We explore a wide range of parameter sets for all models, ensuring consistent parameter ranges for any shared (e.g., extracellular) components.

First, we plot the normalized sensitivities, H_input→output_, which naturally describe the (overall) intracellular signalling feedbacks; if H > 0, then signalling activates the prescribed output, whereas if H < 0, then it inhibits it (Figure 3A). An advantage of considering H is that we can derive strict bounds (light blue lines in Figure 3B-E) that are not only independent of the intracellular dynamics but also apply across all possible extracellular parameters and depend only on the type of inhibitor present (see Supplementary Materials). When we identify pattern-forming parameter sets numerically, we find that, regardless of the intracellular dynamics assumed, the normalized sensitivities all fall within the predicted region i.e., satisfy the general limits defined by **𝒩** (Figure 3B-E). Moreover, we observe that some parameter sets lie close to these limits, demonstrating that the bound is tight. To quantify this further, we directly compare the parameter sets that satisfy **𝒩** with those that form patterns i.e., those that satisfy the necessary and sufficient conditions. Across all types of intracellular dynamics, we find a large overlap (>90%) between the parameter sets that satisfy **𝒩** and the true pattern-forming parameter space (Figure 3B-E); this is also the case when assuming alternative extracellular interactions (Figure S2A-D, Figure S3). We could find parameter regimes where the agreement between **𝒩** and pattern-forming ability was lower (Figure S2E), although this required substantial differences in diffusion rates that may not be relevant *in vivo*. Together, this suggests that, at least for the models and parameter ranges considered here, the general limits that we derive from **𝒩** – which apply regardless of the precise intracellular dynamics – place tight constraints on pattern-forming parameter values and may be reasonably close to being necessary and sufficient conditions for patterning. Furthermore, this implies that key properties of reaction-diffusion systems may be determined by analysing models in which intracellular signalling is approximated by a single effective component, *S*, per pathway and a correspondingly simpler reaction-diffusion matrix 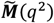.

### Inhibitor type strongly constrains the regulatory logic of pattern-forming activator-inhibitor circuits

So far, we have explored how different *intracellular* dynamics impact the pattern-forming capacity of reaction-diffusion models. Now, we investigate the effect of incorporating different *extracellular* interactions. We restrict our attention to activator-inhibitor systems (i.e., those with just two secreted molecules that either activate or inhibit signalling) and explicitly model the mechanism of pathway inhibition. This leads us to consider four distinct types of (diffusible) inhibitor: (1) inhibitors that bind directly to signalling ligands, sequestering them into an inert complex that eventually degrades (Figure 4A); (2) inhibitors that bind to receptors but do not activate them, thus competing with ligand-induced signalling (Figure 4B); (3) inhibitors that reversibly form complexes with ligands, in which the complex cannot activate signalling but can dissociate to release the ligand (Figure 4C); (4) inhibitors that act indirectly on the activator i.e., ligands that activate other signalling pathways, which then leads to transcriptional inhibition of the activator (Figure 4D). For each inhibitor type, we sought to understand what regulatory interactions were compatible with patterning; for example, the classic activator-inhibitor scheme posits that both activator and inhibitor are induced by signalling (H_s→A_ > 0, H_s→I_ > 0).

**Figure 4.**
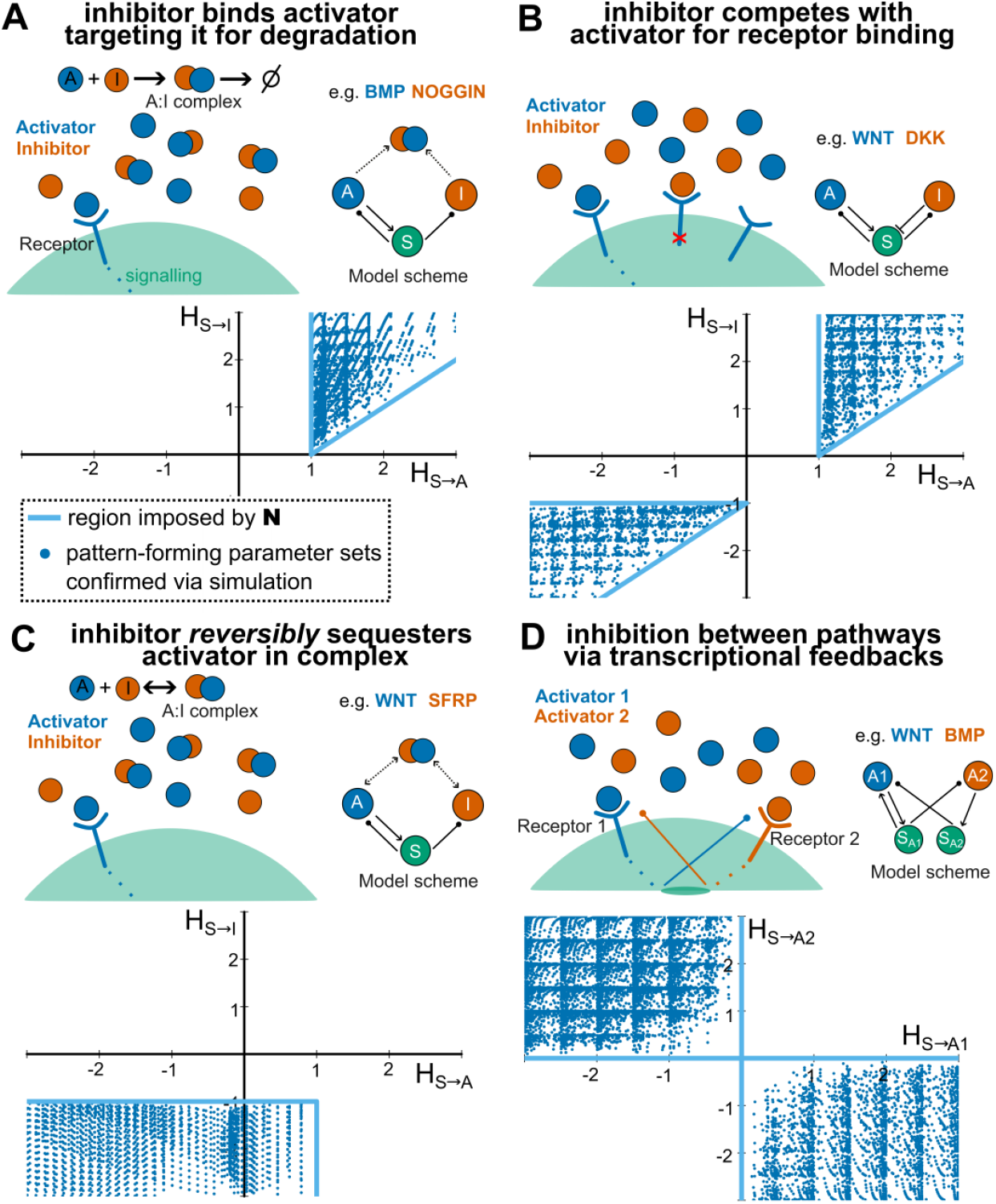
The type of extracellular inhibitor strongly constrains the regulatory logic of pattern-forming activator-inhibitor circuits. (**A-D**) Analysis of activator-inhibitor systems with different types of extracellular interactions (schematics, upper) reveal strict constraints on feedback regulation required for patterning to occur (visualized via the normalized sensitivities H_s→A_, H_s→I_, lower). Shown are analytical limits that apply regardless of the intracellular dynamics (light blue lines), as well as numerical simulations of models in which signalling is approximated by a single variable (blue dots). Parameter ranges were chosen to adequately explore the space defined by the necessary condition **𝒩**. (**A**) When inhibitors bind directly to signalling ligands, sequestering them into inert complexes that eventually degrade, both the activator and the inhibitor must be *activated* by signalling for patterns to form. (**B**) When inhibitors bind to receptors but do not activate them-competing with ligand-induced signalling - the system permits pattern-formation only if signalling either *activates* or *inhibits* the expression of both activator and inhibitor. (**C**) When inhibitors reversibly bind to ligands, forming complexes that cannot activate signalling but can dissociate to release the free ligand, signalling must *inhibit* the expression of the inhibitor. The activator may be *activated, inhibited* or *unaffected* by signalling. (**D**) When inhibitors act indirectly on the activator, i.e., ligands activate other signalling pathways, pattern formation requires one pathway to *activate* expression of its own ligand and exactly one pathway to *inhibit* the ligand of the other.

We take two complementary approaches. First, we derive general analytical limits on the normalized sensitivities, that apply regardless of the assumed intracellular networks and across all intracellular/extracellular parameter values (light blue lines in Figure 4, see Supplementary Materials). Second, we perform numerical simulations assuming a specific intracellular network (for simplicity, a single effective component, *S*, per pathway) and systematically vary parameters to plot a wide range of pattern-forming parameter sets (blue dots in Figure 4, compared with **𝒩** in Figure S3).

Strikingly, we find that each type of diffusible inhibitor requires distinct regulatory circuits to self-organize patterns:

1. When the inhibitor irreversibly sequesters the activator by directly binding to it, then both activator and inhibitor must be *activated* by signalling for patterns to form (Figure 4A); this is consistent with the intuition from classic activator-inhibitor models.
2. When the inhibitor instead competes with the activator for receptor binding, then an additional type of regulation is also compatible with patterning, in which signalling *inhibits* both activator and inhibitor expression (Figure 4B).
3. When the inhibitor binds the ligand, but the resulting complex can readily dissociate, then the classic activator-inhibitor logic (i.e., signalling *activates* both activator and inhibitor) no longer permits patterning. Instead, signalling must always *inhibit* expression of the inhibitor (Figure 4C). The activator may also be *inhibited* by signalling, leading to a similar circuit as above in (2). However, it is also possible that the activator is itself *activated* by signalling, meaning that activator and inhibitor are no longer co-expressed, a scenario not predicted by simpler activator-inhibitor models. Finally, self-organized patterning appears possible even if the activator is uniformly expressed and not impacted by signalling level at all.
4. When inhibition acts indirectly (i.e., when there are two diffusible ligands activating different signalling pathways), we find that one pathway must *activate* expression of its own signalling ligand (see Supplementary Materials); without loss of generality, we define this to be pathway 1. We also require there to be a single *inhibitory* interaction between the pathways (Figure 4D), i.e., activation of pathway 1 *inhibits* the expression of ligand 2 (or vice versa).

We also considered a more general case that combines scenarios (1) and (3), in which the inhibitor and activator form a complex that both degrades *and* dissociates (Figure S4). We find that the circuits able to form patterns are a combination of the limiting cases defined by scenarios (1) and (3), with different circuits possible based on the ratio of the complex dissociation rate to the complex degradation rate (see Supplementary Materials for further discussion).

Together these results demonstrate that, unlike the intracellular components, the extracellular interactions in reaction-diffusion systems strongly impact their pattern-forming potential and lead to qualitatively distinct types of activator-inhibitor circuit. We emphasize that the constraints on the type of regulatory interactions may be derived from the necessary condition **𝒩** alone and thus depend only on the type of diffusible inhibitor present and not on the precise intracellular dynamics assumed.

### An atlas of pattern-forming activator-inhibitor circuits

For each type of inhibitor, and corresponding regulatory circuit(s) identified in Figure 4, we now use numerical simulations via ReactionDiffusion.jl to predict the resulting classes of self-organized patterns (Figure 5). One key difference amongst them is the predicted phase difference between components. Regions of active signalling may be in-phase (i.e., co-expressed) with both the ligand and inhibitor expression domains (Figure 5A, B_1_). Alternatively, signalling may be out-of-phase (i.e., anticorrelated) with both ligand and inhibitor expression (Figure 5B_2_,C_3_). Beyond these two scenarios, the ligand need not always be co-expressed with the inhibitor but may instead be expressed in regions where inhibitor levels are low (Figure 5C_1_) or present uniformly throughout the tissue (Figure 5C_2_). Finally, when two distinct signalling pathways are involved, the two ligands may either be co-expressed in the same location (Figure 5D_1_) or anticorrelated and thus expressed in anti-phase to each other (Figure 5D_2_). These phase differences provide concrete, qualitative predictions – independent of specific parameter values – that may then be compared to measurements of gene expression patterns *in vivo*.

**Figure 5.**
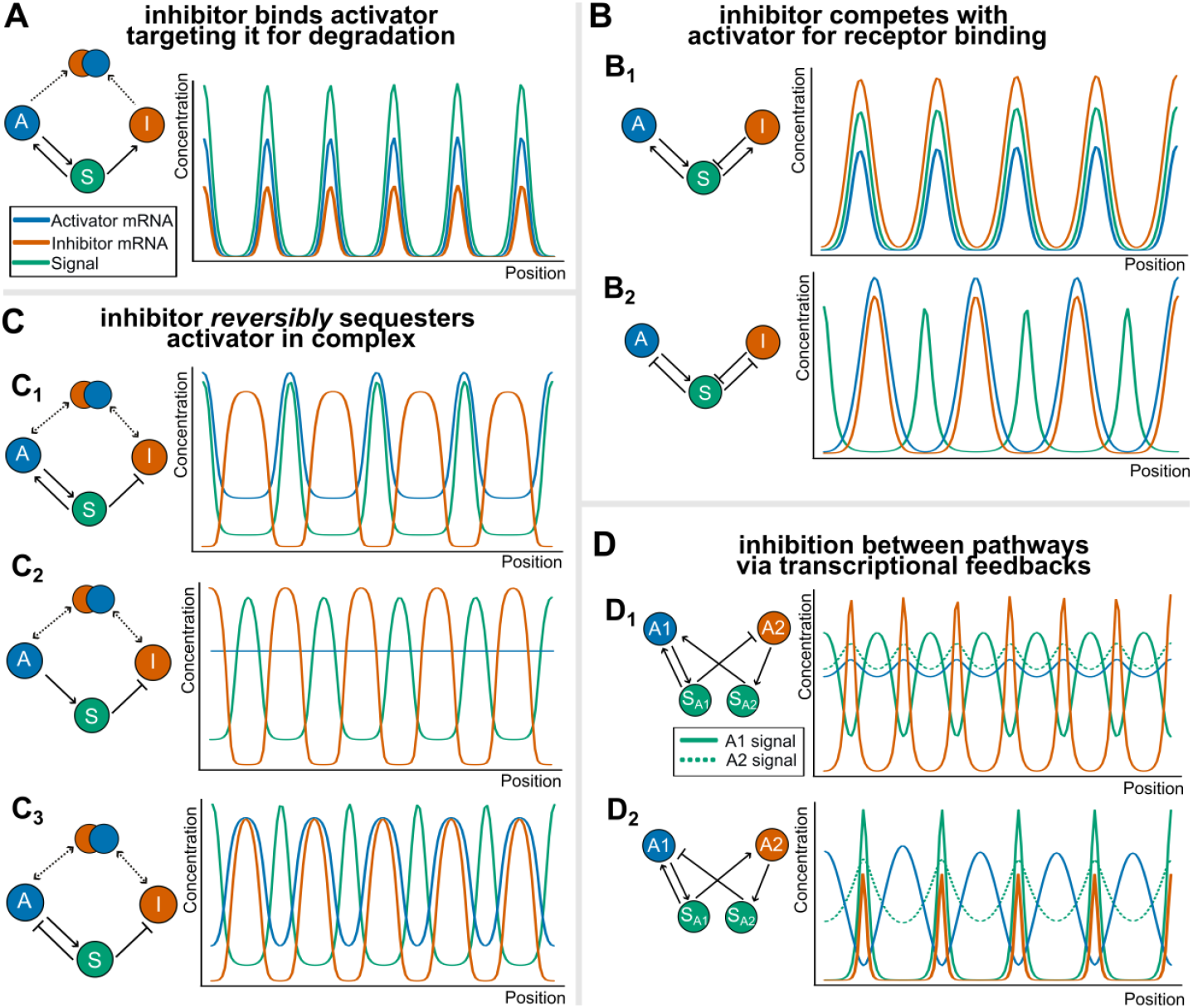
An atlas of pattern-forming activator-inhibitor circuits. (**A-D**) Predicted expression patterns for each type of inhibitory mechanism and corresponding regulatory circuit(s) from Figure 4. Plots show spatial distributions of relative mRNA concentrations for the activator (blue) and inhibitor (red), alongside levels of signalling pathway activation (green). (**A**) Pathway activation is co-expressed (in-phase) with both the activator and inhibitor expression domains. (**B**) Pathway activation is either in-phase (**B**_**1**_) or out-of-phase (**B**_**2**_) with both the activator and the inhibitor. (**C**) Pathway activation is out-of-phase (anticorrelated) with the inhibitor, and is either: in-phase with the activator (**C**_**1**_), uniform across the tissue (**C**_**2**_), or out-of-phase with the activator (**C**_**3**_). (**D**) The two ligands of the two signalling pathways are either co-expressed in the same location (**D**_**1**_) or expressed out-of-phase with each other (**D**_**2**_).

We have also begun to explore the parameter-dependence of pattern formation for each of the circuits. We have found, and show with examples in Figure S5, that there exist parameter values that are predicted to form self-organized patterns in a biologically plausible range. Moreover, for these representative parameter sets, we show that patterning can still occur despite significant variation in any of the individual parameters of the models (Figure S5, left column). This suggests that, at least for the regions of parameter space shown here, the ability of activator-inhibitor circuits to form patterns can be highly robust to parameters. We also computed robustness metrics for each parameter individually by calculating the fraction of parameter space that forms patterns (Figure S5, right column). This reveals general tendencies that affect the ability of each of the circuits to form patterns; for example, we predict that a highly diffusive complex of activator and inhibitor increases the likelihood of the circuit being pattern forming (Figure S5C). We find that these broad tendencies can be derived by considering the necessary condition **𝒩** alone and are therefore likely to apply regardless of the precise intracellular dynamics present. Together, these results point to eight distinct classes of pattern that can self-organize via activator-inhibitor mechanisms.

## DISCUSSION

Activator-inhibitor circuits have been widely hypothesized to self-organize patterns in a variety of developing tissues (Kondo and Miura, 2010; Marcon and Sharpe, 2012). Here, we develop activator-inhibitor models that are more closely aligned to *in vivo* signalling mechanisms and find that they display qualitatively distinct behaviours to simpler, phenomenological approximations. In particular, the mode of extracellular inhibition strongly constrains the type of regulatory architectures capable of self-organizing patterns *de novo*. In contrast, the ability to break symmetry is relatively insensitive to the intracellular dynamics and depends only on the overall, steady-state response of the cell to signalling, which can often be approximated or estimated directly. We have combined formal mathematical proofs with large scale numerical simulations to support the generality of our conclusions, and derive strict constraints on patterning (via the necessary condition **𝒩**) which apply regardless of intracellular signalling mechanisms. These constraints do not guarantee that a pattern will form, nor do they predict all features of the resulting patterns (e.g., spatiotemporal dynamics, final pattern formed), but they do point to broadly applicable principles that are necessary for reaction-diffusion systems to self-organize.

Guided by these principles, we systematically characterize systems that can form patterns with only a single activator and a single inhibitor, and identify key gene expression signatures associated with each class of mechanism. These include the classic Turing-type logic, in which signalling promotes the simultaneous expression of both activator and inhibitor. However, several other circuits form patterns but have an inverse relationship between signalling and activator/inhibitor production. Indeed, recent experimental observations point to numerous examples of this inverted logic operating *in vivo* (Grall et al., 2024; Grocott et al., 2020). When we allow the inhibitor to *reversibly* bind the signalling ligand, we predict several further types of circuit capable of breaking symmetry. This includes cases where activator and inhibitor are expressed in opposite locations, a property not predicted by simpler activator-inhibitor models, but something that has been observed for several self-organizing patterns *in vivo* (e.g., WNT/sFRP in cortical organoids (Takata et al., 2017)). Moreover, we predict that an elevated diffusivity of the ligand in complex promotes pattern formation (Figure S5C), suggesting that ligand shuttling (as has been suggested for e.g., WNT/sFRP (Mii and Taira, 2011, 2009), BMP-Chordin (Zhu et al., 2023)) may also be important (Shilo et al., 2013). Together, our comprehensive atlas of activator-inhibitor mechanisms reveals a more varied repertoire of pattern-forming motifs than previous recognized which may be operating across a diverse range of tissues *in vivo*.

When exploring the parameter-dependence of our models, we find instances where patterning is both robust to parameter variation and occurs within a biologically feasible range, hinting that systems consisting of only a single activator/inhibitor pair might, by themselves, be capable of highly robust patterning. However, our analysis of model robustness is rather preliminary, characterizing either isolated exemplar parameter sets (Figure S5, left column) or collapsing the full parameter space down to one dimension for ease of visualization (Figure S5, right column). In future, more work will be needed to fully characterize the high-dimensional parameter spaces and thus robustness of each of the models in Figure 5, using tools and approaches applied elsewhere. Moreover, integrating model predictions with *in vivo* data and biophysical measurements of key parameters will also be necessary to definitively test the hypothetical circuits that we have proposed here (Kuhn et al., 2022; Müller et al., 2012).

A further limitation of this study is that we restrict our attention to mechanisms with only two secreted diffusible molecules. Whilst some *in vivo* patterns do appear to involve single activator/inhibitor pairs (e.g., GDF5/NOG in the developing finger joints (Grall et al., 2024)), others are suggested to involve multiple signals and pathways (Economou et al., 2021; Glover et al., 2017; Raspopovic et al., 2014). Moreover, *in silico* studies have shown that adding diffusible molecules not only expands the number of possible pattern-forming networks but can also be associated with increased robustness (Marcon et al., 2016; Scholes et al., 2019; Shaberi et al., 2025). In addition, our models have so far neglected impacts of receptor dynamics on extracellular ligands and inhibitors (e.g., receptor-mediated ligand sequestration (Lord et al., 2019; Preiß et al., 2022), endocytic ligand recycling (Romanova-Michaelides et al., 2022)), and have focused on activator-inhibitor complexes with 1:1 stoichiometries (like GDF5-NOG). Despite these simplifications, our models capture the minimal mechanistic features associated with intercellular signalling and distinct types of pathway inhibition, and this already revealed qualitative differences with phenomenological Turing models. We expect that incorporating more *in vivo* interactions and complexity into our models will further expand our understanding of reaction-diffusion based patterning.

Beyond the principles we have uncovered, the computational frameworks that we have developed are broadly applicable to any reaction-diffusion mechanism. We have built an end-to-end, freely-available simulation pipeline (ReactionDiffusion.jl) that generalizes to mechanisms with any number of components/interactions and that may be used without detailed knowledge or fine tuning of PDE solvers. We are currently limited to one dimensional simulations, which are sufficient to determine the overall characteristics of a pattern (i.e., can it form *de novo?* what components are co-expressed?) but can often not be compared directly to experimental data. Moreover, we have restricted our attention to fully self-organizing systems (i.e., no prior asymmetries in the tissue) with a strong focus on periodic patterns. Future work will be required to extend our pipeline to broader classes of pattern with more realistic geometries and external/initial asymmetries.

Together, this work has revealed strong constraints and general design principles for self-organized patterning via reaction-diffusion mechanisms. The candidate circuits we have identified may point experimentalists to new activator-inhibitor circuits that may be present *in vivo*, and also suggest promising designs for synthetic morphogen circuits that can self-organize *in vitro* (Sekine et al., 2018; Tica et al., 2024; Toda et al., 2020). Combined with experimental advances in spatiotemporal measurement and perturbation, we are now poised to further understand the implications and implementation of Turing’s reaction-diffusion hypothesis *in vivo*.

## Code availability

We have developed an open-source Julia package with extensive documentation and tutorials to simulate complex reaction-diffusion systems: https://github.com/hiscocklab/ReactionDiffusion.jl.

We provide a separate repository for the code used to generate the results in this paper: https://zenodo.org/records/15495853

## Acknowledgements

We would like to thank members of the Hoppler Lab, and Aberdeen Developmental Biology Group for their useful feedback throughout the project. This work has been supported by ERC grant SELFORG-101161207, and UK Research and Innovation (Biotechnology and Biological Sciences Research Council, grant number BB/W003619/1).

## Funding disclaimer

Funded by the European Union. Views and opinions expressed are however those of the author(s) only and do not necessarily reflect those of the European Union or the European Research Council Executive Agency. Neither the European Union nor the granting authority can be held responsible for them

---

**Figure S1.**
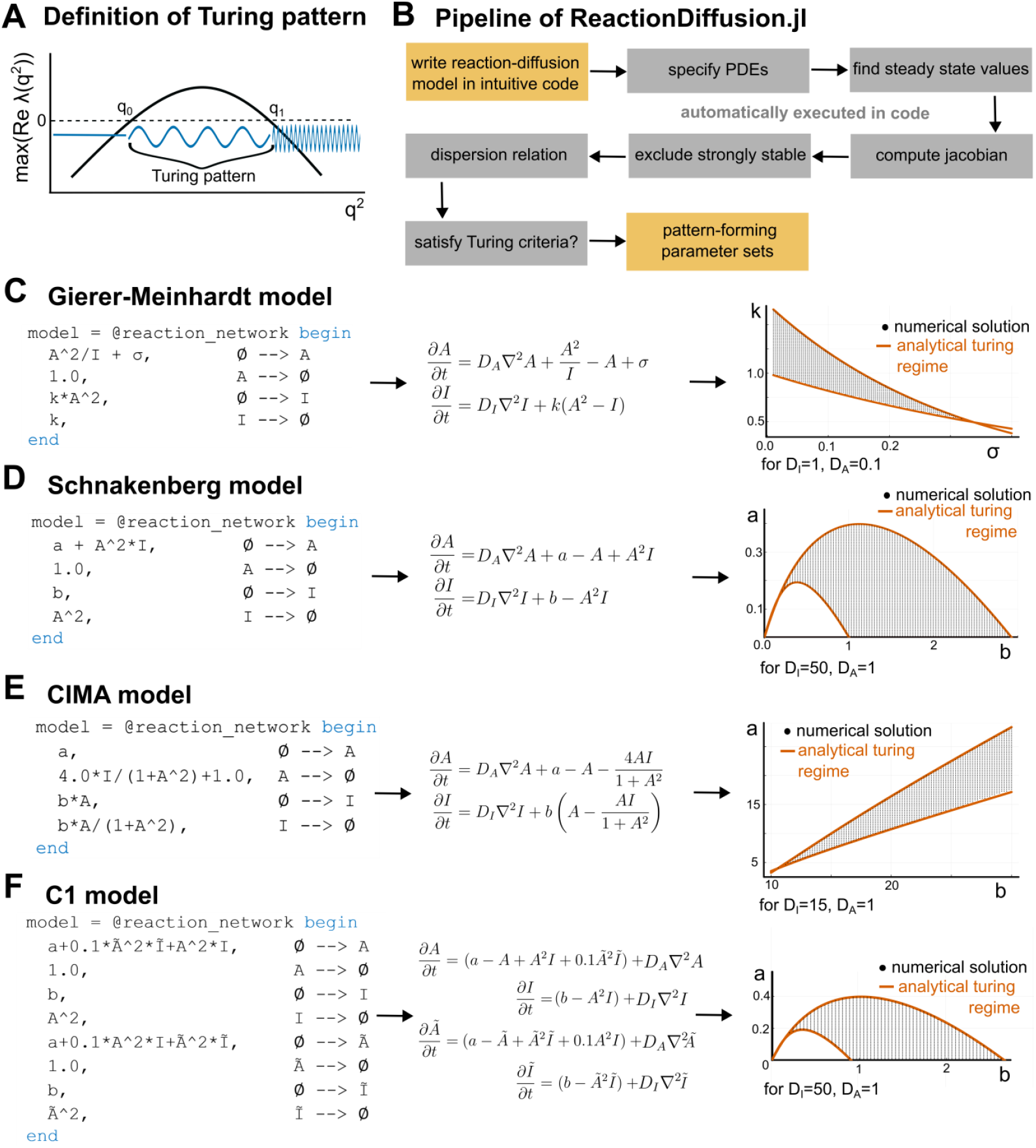
Detection of Turing instabilities in ReactionDiffusion.jl. (**A**) Schematic definition of a (Type I) Turing instability, where λ(q^2^) refers to the eigenvalues of the linearized reaction-diffusion matrix as a function of pattern wavevector. (**B**) Outline of computational pipeline that automates the identification of pattern-forming parameter sets in ReactionDiffusion.jl. (**C-F**) Benchmarking our detection of Turing instabilities against previously studied Turing systems (**C**) Gierer-Meinhardt (Meinhardt and Gierer, 1974), (**D**) Schnakenberg (Schnakenberg, 1979), (**E**) CIMA (Lengyel and Epstein, 1991), (**F**) 4-component, coupled Schnakeberg (Kurics et al., 2014)). Each previous model is represented in intuitive code (left). This is automatically converted to a set of PDEs (middle). We plot parameter combinations that we predict undergo a Turing instability (black dots) and find that these agree closely with analytically derived limits in all cases (red lines, see Appendix 1).

**Figure S2.**
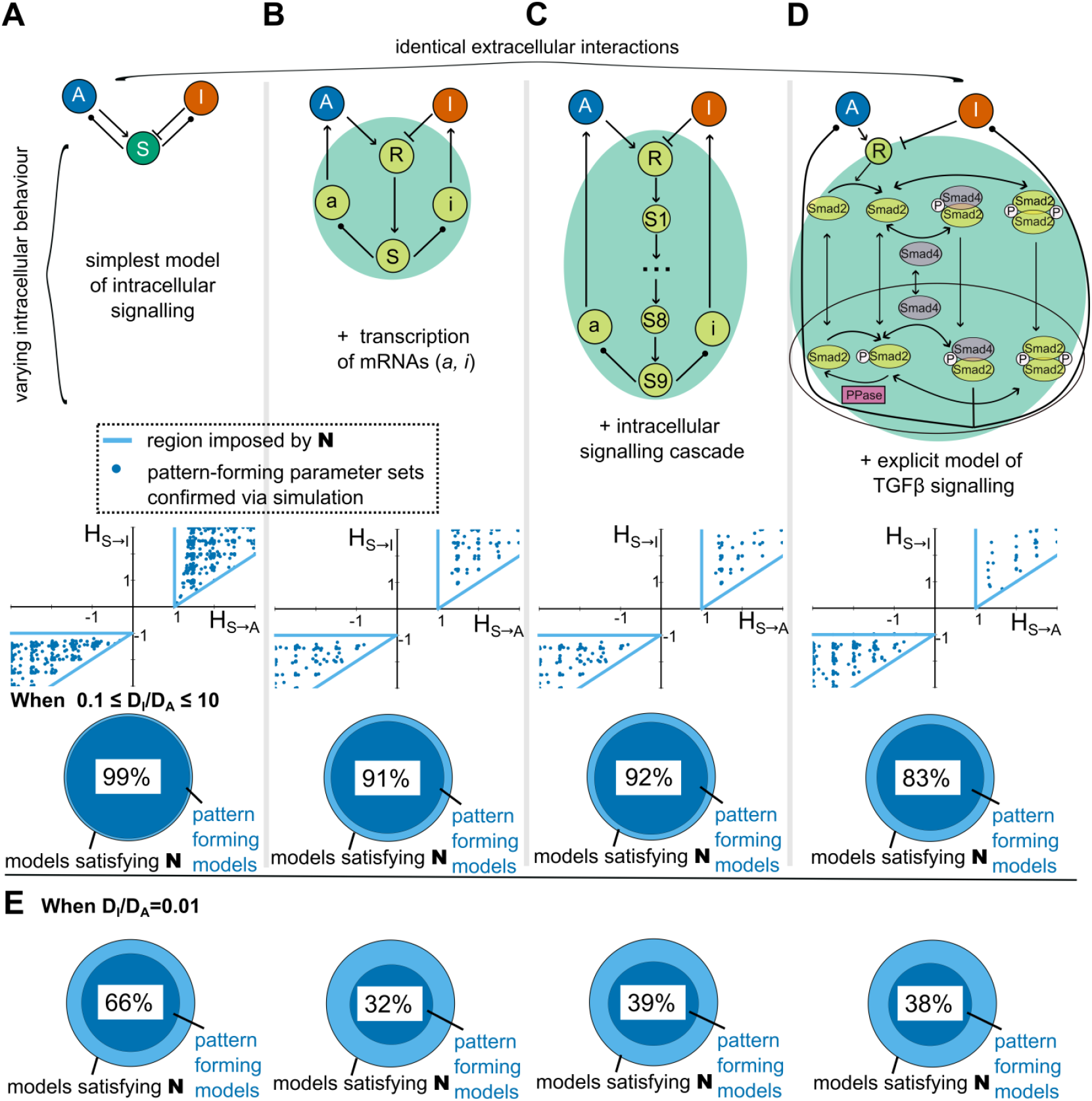
Exploring the effect of intracellular dynamics in models where the inhibitor competes with the activator for receptor binding. (**A-D**) Analysis of activator-inhibitor systems with the same extracellular but different intracellular interactions. In contrast to Figure 2, here we assume that the extracellular inhibitor competes with the activator for receptor binding. Across a variety of intracellular behaviours (schematics, upper row), we find that the sensitivity parameters are subject to the same constraints across all models (light blue lines, middle row), which we confirm via numerical simulation (blue dots, middle row). As in Figure 2, we find that the necessary condition **𝒩** alone can reasonably approximate the full set of pattern-forming parameters (~80% agreement, lower row). (**E**) Example parameter regime in which the necessary condition **𝒩** does not provide a good approximation of the pattern-forming parameter space. For this model, when the diffusion ratio is small,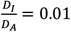 = 0.01, the agreement between the parameters satisfying **𝒩** and the parameters that permit patterning drops below 40%.

**Figure S3.**
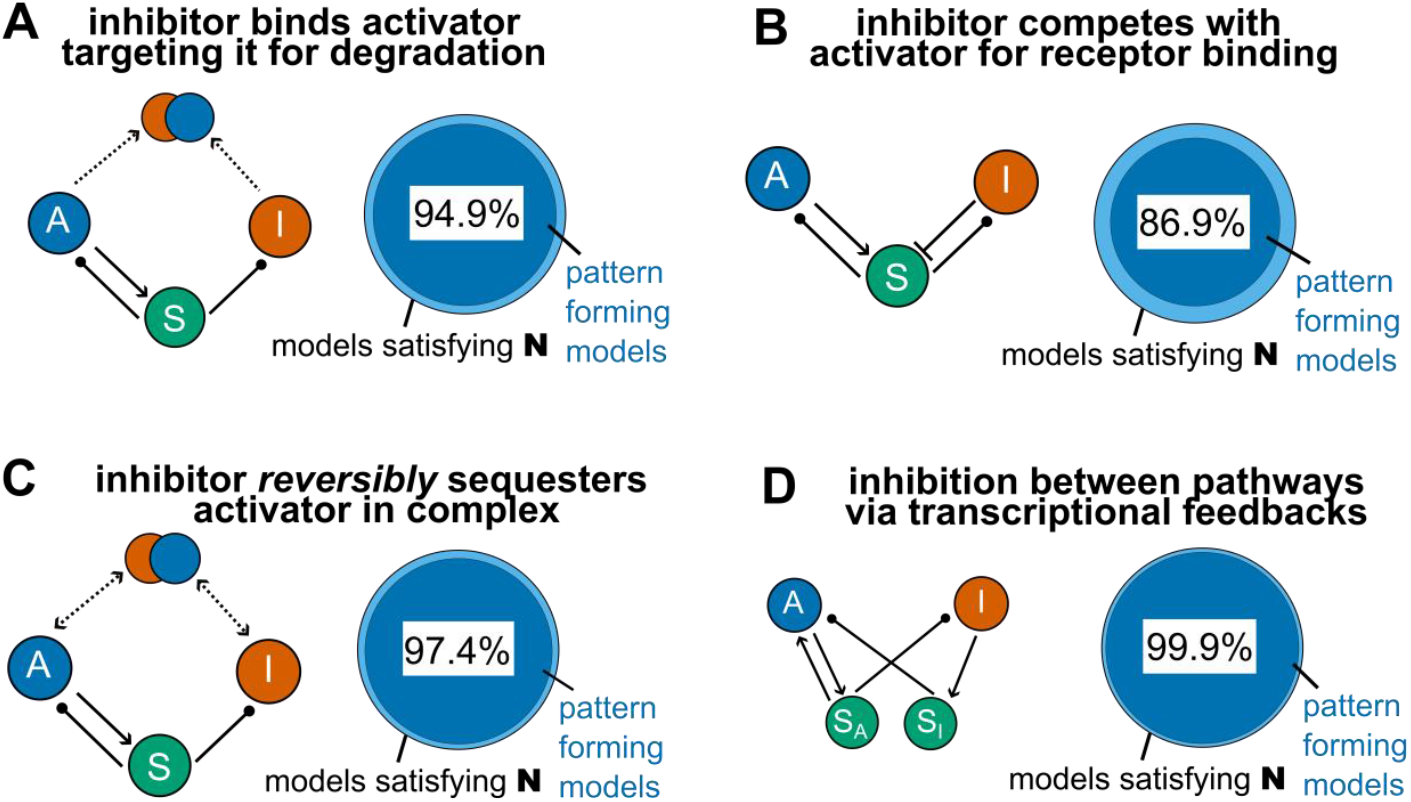
Pattern-forming parameters of the simplified models can be reasonably approximated by the necessary condition 𝒩 alone. (**A-D**) Further analysis of the models from Figure 4. The pattern-forming parameter sets are compared to the parameter sets satisfying the necessary condition **𝒩** alone. We find an agreement of at least 86% for all models.

**Figure S4.**
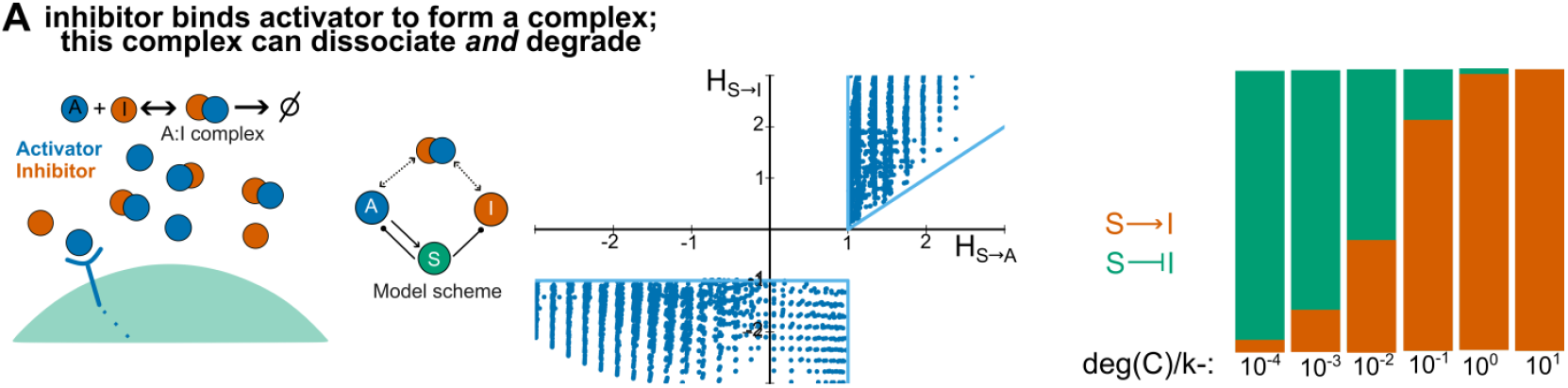
Models in which the activator and inhibitor form a complex that cannot signal, but can both dissociate and degrade. (A) Analysis of the case in which the activator-inhibitor complex both dissociates and degrades. As expected, the predicted regulatory interactions (H_s→A_, H_s→I_), span both limiting cases, in which the complex degrades but does not dissociate (Figure 4A), or dissociates but does not degrade (Figure 4C). Consistent with our analysis (see Supplementary Materials), numerical simulations confirm that the ratio of complex degradation to dissociation rates dictates which regulatory circuit(s) are possible (right).

**Figure S5.**
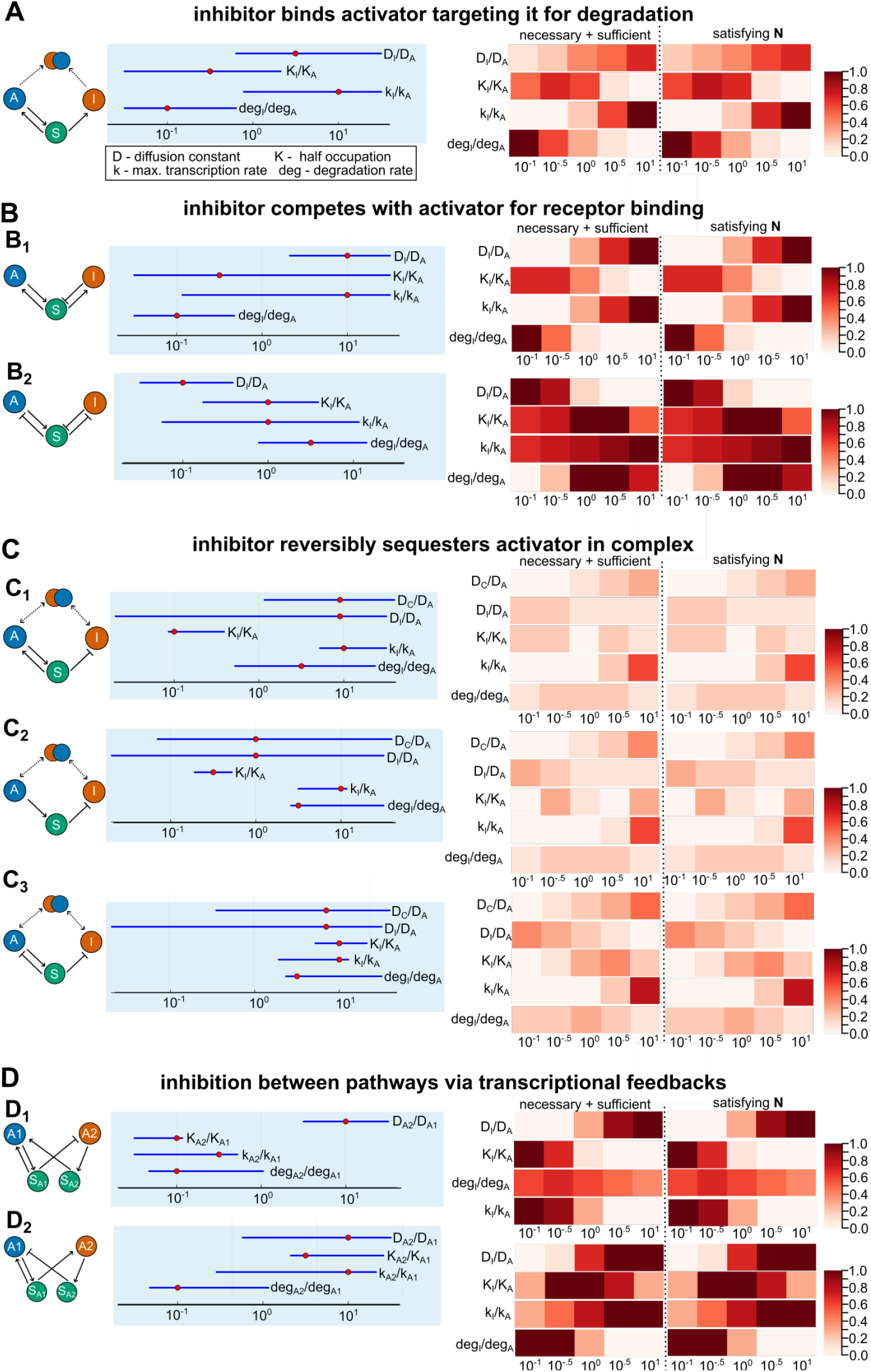
Robustness of pattern-forming activator-inhibitor circuits. (**A-D**) Exploration of parametric robustness for each type of activator-inhibitor circuit shown in Figure 5, showing two different robustness measures. Left: Starting from the parameter values used in Figure 5 (red dots), we show how much each parameter can be individually varied (blue lines) without compromising the ability of the model to form patterns. Right: Heatmaps denoting the fraction of parameter sets (colorbar, measured in %) that are predicted to break symmetry. Each row corresponds to fixing a different parameter, with all other parameters allowed to vary across a wide range (0.1 to 10). We find that this robustness metric aligns closely with the prediction based solely on the necessary condition **𝒩**.

