## Supplementary Materials for "Fundamental limits on patterning by Turing-like reaction-diffusion mechanisms"

### Supplementary Material

**3 Analysis of activator-inhibitor models** **9**
3.1 Case 1: The inhibitor binds to the ligand, inactivating it and targeting it for degradation . . 9
3.2 Case 2: The inhibitor binds to the receptor, competing with the ligand for pathway activation 11

### 1 *ReactionDiffusion.jl*: a computational pipeline to simulate complex reaction-diffusion PDEs

We have developed a pipeline to simulate arbitrary reaction-diffusion systems using the performant and mature SciML ecosystem in Julia. We restricted ourselves to a simple, one-dimensional geometry and focused exclusively on reaction-diffusion PDEs (i.e., those in which the only spatial derivative terms are Laplacian).

The reaction terms are specified in an intuitive domain specific language that naturally captures biochemical interactions, as developed in the *Catalyst.jl* package [1]. Here, reaction rates are either prescribed by mass action kinetics (i.e., determined by the stoichiometry of the interactions) or, where applicable, by user-defined functions. We approximate the diffusion operators via a finite difference method (with reflective boundary conditions) in one dimension, which, via the method of lines, converts the system of PDEs into a large set of ODEs operating on a discretized grid.

The resulting ODEs are solved using the *ARK4(3)6L[2]SA* algorithm (KenCarp4 in *DifferentialEquations.jl*) (see [2]), an A-L stable, stiffly-accurate 4th order ESDIRK method which we found to be highly effective for the stiff PDEs associated with reaction-diffusion systems. The simulation time was significantly reduced by using *Symbolics.jl* and *ModelingToolkit.jl* to symbolically compute the very large, but sparse Jacobian terms; previous work has shown that this can accelerate implicit PDE solvers by several orders of magnitude [3].

We use adaptive time-stepping (via KenCarp4 with tolerances: relative=1e-6, absolute =1e-8) and terminate the simulations once they have reached a steady state, meaning that the step-size/timestep and end point of the simulations need not be known or specified a priori and is instead automatically adjusted throughout the simulation.

We have put all these components together inside a single, easy-to-use function, that serves as a fully end-to-end pipeline to simulate reaction-diffusion PDEs of arbitrary complexity. Models can be built quickly in a few lines of code and simulated in a computationally efficient manner, without specifying solver type or fine tuning algorithm parameters. We make this pipeline freely available as a Julia package, *ReactionDiffusion.jl*, with documentation, tutorials and examples provided: https://github.com/hiscocklab/ReactionDiffusion.jl.

#### 1.1 Automated detection of Turing instabilities in *ReactionDiffusion.jl*

A necessary condition for a Turing pattern to form is that diffusion causes a homogeneous steady state to become unstable with respect to periodic disturbances of certain wavelengths. We implement standard techniques from linear instability analysis into *ReactionDiffusion.jl* to automatically detect whether a given reaction-diffusion PDE undergoes such a diffusion-driven (i.e., Turing) instability.

First, we neglect diffusion and use an ODE solver to find the homogeneous steady state of the system. (For the models considered here, there is typically a single stable steady state; in general, there may be multiple stable states, which can be found by varying the initial conditions of the ODE simulation in *ReactionDiffusion.jl*, following the approach of [4]). We use a fifth order Rosenbrock method (Rodas5 in *DifferentialEquations.jl*) to efficiently solve the ODEs until they reach a steady state. We then compute the Jacobian of the reaction terms about this steady state to arrive at the reaction component, ($\mathbf{J}$), of the linearized reaction-diffusion matrix.

Following [5] and [6], we determine whether all signed principal minors of $\mathbf{J}$ are non-negative, in which case it has been shown the system cannot undergo a Turing instability regardless of the diffusion parameters. We also check that $\mathbf{J}$ is stable (i.e., all eigenvalues have negative real part), which is to be expected since we have simulated the long-time behaviour of the ODEs to compute the stationary point. Then, following the approach of [4], we compute the eigenvalues of $\mathbf{J} - q^2\mathbf{D}$ for a wide range of values for the wavevector $q$ (log-varying over a 10,000 range). We restrict our attention to Type I Turing instabilities, defined as those for which there exists an eigenvalue of $\mathbf{J} - q^2\mathbf{D}$ with positive real part for some finite range of (positive) wavevectors. Together, these steps determine whether a given set of reaction-diffusion PDEs, for a given set of parameters, undergoes a (type I) Turing instability.

We repeat this analysis in parallel to systematically screen all combinations of user-provided parameter values, using multithreading (of both the ODE solver and the eigenvalue calculation steps) to significantly accelerate the large-scale parameter screens. All of these steps are combined into a single, easy-to-use function, which uses the same intuitive model syntax as described above. Together, this allows users to detect Turing instabilities in reaction-diffusion PDEs with just a few lines of code and for millions of parameter sets per minute on a standard desktop.

To evaluate the accuracy of our approach, we applied our pipeline to several well-studied Turing systems for which we can derive analytical expressions for the parameter constraints required for Turing instabilities. We have investigated three classic Turing systems: Schnakenberg [7], CIMA [8], Gierer-Meinhardt [9], as well as a more recent 4-component system [10]. In all cases, we find excellent agreement between our numerical predictions and the analytical ground truth. See Appendix 1 for a detailed derivation of the analytical conditions.

### 2 Deriving necessary conditions for reaction-diffusion systems to break symmetry

#### 2.1 The necessary condition, $\mathcal{N}$, must hold for a Turing-instability to occur

We model intercellular signalling by the following system of reaction-diffusion partial differential equations (PDEs):

$$\frac{\partial \mathbf{c}}{\partial t} = \mathbf{f}(\mathbf{c}) + \mathbf{D}\nabla^2 \mathbf{c}, \tag{1}$$

where $\mathbf{c}$ is a vector of all species of given system, $\mathbf{f} : \mathbb{R}^n \to \mathbb{R}^n$ is a set of nonlinear functions denoting the interactions between individual species, $\mathbf{D}$ is a diagonal matrix with diffusion rates on the diagonal, and the $\nabla^2$ operator stands for divergence of the gradient, i.e., Laplacian.

Let $\mathbf{M}(q^2) \stackrel{\text{def}}{=\!=} \mathbf{J} - q^2\mathbf{D}$ be the reaction-diffusion matrix, linearised about the homogeneous steady state defined by $\mathbf{f}(\bar{\mathbf{c}}) = \mathbf{0}$, where $\mathbf{J}$ is the Jacobian of the reaction terms, and $q$ is the wavenumber corresponding to wavelength $\lambda = 2\pi/q$. A necessary condition for a Turing instability is that there exists a stable homogeneous steady state (in the absence of diffusion) which becomes unstable for some finite values of wavevector $q$. To be precise, we say the system is pattern-forming if there exist two positive real numbers $q_0, q_1$ such that the system is stable (i.e., all eigenvalues have negative real parts) on the interval $[0, q_0) \cup (q_1, \infty)$ and has exactly one positive real eigenvalue on the interval $(q_0, q_1)$ (see Figure S1-A). This strict definition excludes some other systems capable of forming patterns [11], but is used here to keep the discussion mathematically rigorous.

For a system consisting of $n$ species, $\mathbf{M}$ is a $n \times n$ matrix. The determinant is equal to the product of its eigenvalues: $det(\mathbf{J} - q^2\mathbf{D}) = |\mathbf{J} - q^2\mathbf{D}| = \prod_{i=1}^{n} \lambda_i$. When the system is stable, all eigenvalues have negative real parts, and so the signed product of eigenvalues must be positive, i.e., $(-1)^n|\mathbf{J} - q^2\mathbf{D}| > 0$. When the system exhibits Turing instability, exactly one real eigenvalue turns positive and therefore $(-1)^n|\mathbf{J} - q^2\mathbf{D}|$ turns negative (Diego et al., 2018). Therefore, $(-1)^n|\mathbf{J} - q^2\mathbf{D}|$ strictly negative for $q \in (q_0, q_1)$ and strictly positive for all other $q$ are necessary conditions for a system to be pattern forming. This requires $|\mathbf{J} - q^2\mathbf{D}|$ to change sign twice for positive $q$, a fact we hereafter refer to as the necessary condition $\mathcal{N}$.

This condition is not sufficient, because it does not rule out that the Jacobian $\mathbf{J}$ itself is unstable or that other eigenvalues change sign as well.

#### 2.2 Simplification of the necessary condition $\mathcal{N}$ applied to models of developmental signalling pathways

In general, signalling pathways comprise elements which diffuse between cells (e.g., extracellular ligands and inhibitors; we label these $D$); elements which directly interact/bind to the diffusible elements (e.g. receptors; we label these as $R$); and elements which are confined within the cell itself (e.g., intracellular signal transduction elements; we label these as $I$).

The matrix $\mathbf{J} - q^2\mathbf{D}$ can be partitioned accordingly into submatrices that represent the interactions between any two groups. We denote by $\mathbf{J}_{ij}$ the block of weights of reactions from $j$ to $i$. For example, $\mathbf{J}_{RD}$ contains all interactions in which diffusible extracellular species affect receptors; $\mathbf{J}_{II}$ contains all interactions between intracellular elements. Likewise, $\mathbf{D}_{DD}$ is a diagonal submatrix containing the diffusion coefficients of the extracellular diffusible species.

Performing this partitioning, we may write:

$$\mathbf{M}(q^2) = \mathbf{J} - q^2\mathbf{D} = \begin{pmatrix} \mathbf{J}_{DD} - q^2\mathbf{D}_{DD} & \mathbf{J}_{DR} & \mathbf{J}_{DI} \\ \mathbf{J}_{RD} & \mathbf{J}_{RR} - q^2\mathbf{D}_{RR} & \mathbf{J}_{RI} \\ \mathbf{J}_{ID} & \mathbf{J}_{IR} & \mathbf{J}_{II} - q^2\mathbf{D}_{II} \end{pmatrix}. \tag{2}$$

This may be simplified by using the fact that the intracellular components, as well as the receptors, do not contribute to intercellular diffusion, and so $\mathbf{D}_{II} = \mathbf{0}$ and $\mathbf{D}_{RR} = \mathbf{0}$. The intracellular components are also not directly affected by the intracellular species, and so $\mathbf{J}_{ID} = \mathbf{0}$. Therefore, for models of signalling pathways it holds that

$$\mathbf{M}(q^2) = \begin{pmatrix} \mathbf{J}_{DD} - q^2\mathbf{D}_{DD} & \mathbf{J}_{DR} & \mathbf{J}_{DI} \\ \mathbf{J}_{RD} & \mathbf{J}_{RR} & \mathbf{J}_{RI} \\ \mathbf{0} & \mathbf{J}_{IR} & \mathbf{J}_{II} \end{pmatrix}.$$

Further, it may often be the case that receptors are not directly affected by intracellular species and therefore $\mathbf{J}_{RI} = \mathbf{0}$. However, to keep our discussion more general, we allow $\mathbf{J}_{RI}$ to be non-zero.

By the Schur complement (see for example [12])

$$det(\mathbf{M}(q^2)) = \begin{vmatrix} \mathbf{J}_{DD} - q^2\mathbf{D}_{DD} & \mathbf{J}_{DR} & \mathbf{J}_{DI} \\ \mathbf{J}_{RD} & \mathbf{J}_{RR} & \mathbf{J}_{RI} \\ \mathbf{0} & \mathbf{J}_{IR} & \mathbf{J}_{II} \end{vmatrix} = |\mathbf{J}_{II}| \cdot \begin{vmatrix} \mathbf{J}_{DD} - q^2\mathbf{D}_{DD} & \mathbf{J}_{DR} - \mathbf{J}_{DI}\mathbf{J}_{II}^{-1}\mathbf{J}_{IR} \\ \mathbf{J}_{RD} & \mathbf{J}_{RR} - \mathbf{J}_{RI}\mathbf{J}_{II}^{-1}\mathbf{J}_{IR} \end{vmatrix} = |\mathbf{J}_{II}| \cdot det(\widetilde{\mathbf{M}}(q^2)), \tag{3}$$

where $|\mathbf{J}_{II}|$ is some real constant and $\widetilde{\mathbf{M}}(q^2)$ is the linearized reaction-diffusion matrix of a simpler model, which we hereafter refer to as the *reduced* model.

This means that,

$$\mathbf{M} = \begin{vmatrix} \mathbf{J}_{DD} - q^2\mathbf{D}_{DD} & \mathbf{J}_{DR} & \mathbf{J}_{DI} \\ \mathbf{J}_{RD} & \mathbf{J}_{RR} & \mathbf{J}_{RI} \\ \mathbf{0} & \mathbf{J}_{IR} & \mathbf{J}_{II} \end{vmatrix} \text{ and } \widetilde{\mathbf{M}} = \begin{vmatrix} \mathbf{J}_{DD} - q^2\mathbf{D}_{DD} & \mathbf{J}_{DR} - \mathbf{J}_{DI}\mathbf{J}_{II}^{-1}\mathbf{J}_{IR} \\ \mathbf{J}_{RD} & \mathbf{J}_{RR} - \mathbf{J}_{RI}\mathbf{J}_{II}^{-1}\mathbf{J}_{IR} \end{vmatrix}$$

are two polynomials that change signs simultaneously for the same values of $q$. This implies that the necessary condition $\boldsymbol{\mathcal{N}}$ for the full linearized reaction-diffusion matrix $\mathbf{M}$ may be derived by considering the reduced matrix $\widetilde{\mathbf{M}}$ alone. Now, by itself, this does not necessarily simplify the calculation of $\boldsymbol{\mathcal{N}}$, since the computation of $\widetilde{\mathbf{M}}$ would require manipulation of the intracellular parts of the Jacobian (e.g., computing $\mathbf{J}_{II}^{-1}$). However, in the next section we show how $\widetilde{\mathbf{M}}$ may be computed more straightforwardly without detailed knowledge of $\mathbf{J}_{II}$.

The equivalence of $\boldsymbol{\mathcal{N}}$ for $\mathbf{M}$ and $\widetilde{\mathbf{M}}$ makes another prediction, namely that the range of possible pattern wavevectors (i.e., those that can destabilize an initially homogeneous state) are the same for $\mathbf{M}$ and $\widetilde{\mathbf{M}}$. This means that we can estimate allowed pattern wavelengths from considering $\widetilde{\mathbf{M}}$ alone. Furthermore, as we shall now show, $\widetilde{\mathbf{M}}$ may also be used to deduce the relative phases between components during symmetry breaking i.e., it can predict which species will be in- or out-of-phase with one another.

###### 2.2.1 Predicting resulting pattern from the reduced system $\widetilde{\mathbf{M}}$

Let $\lambda_q$ be the real eigenvalue that turns positive at $q$ and let $\mathbf{v} = \begin{pmatrix} \mathbf{v}_D & \mathbf{v}_R & \mathbf{v}_I \end{pmatrix}^T$ be the corresponding $n \times 1$ eigenvector. Then it holds that

$$\begin{pmatrix} \mathbf{J}_{DD} - q^2\mathbf{D}_{DD} & \mathbf{J}_{DR} & \mathbf{J}_{DI} \\ \mathbf{J}_{RD} & \mathbf{J}_{RR} & \mathbf{J}_{RI} \\ \mathbf{0} & \mathbf{J}_{IR} & \mathbf{J}_{II} \end{pmatrix} \cdot \begin{pmatrix} \mathbf{v}_D \\ \mathbf{v}_R \\ \mathbf{v}_I \end{pmatrix} = \lambda_q \cdot \begin{pmatrix} \mathbf{v}_D \\ \mathbf{v}_R \\ \mathbf{v}_I \end{pmatrix}. \tag{4}$$

By expressing $\mathbf{v}_I$ as $\mathbf{J}_{DI}(\lambda_q \mathbf{I} - \mathbf{J}_{II})^{-1}\mathbf{J}_{IR}$, we obtain an equivalent system of two equations

$$(\mathbf{J}_{DD} - q^2\mathbf{D}_{DD}) \cdot \mathbf{v}_D + (\mathbf{J}_{DR} + \mathbf{J}_{DI} \cdot (\lambda_q \mathbf{I} - \mathbf{J}_{II})^{-1} \cdot \mathbf{J}_{IR}) \cdot \mathbf{v}_R = \lambda_q \cdot \mathbf{v}_D,$$

$$\mathbf{J}_{RD} \cdot \mathbf{v}_D + (\mathbf{J}_{RR} + \mathbf{J}_{RI} \cdot (\lambda_q \mathbf{I} - \mathbf{J}_{II})^{-1} \cdot \mathbf{J}_{IR}) \cdot \mathbf{v}_R = \lambda_q \cdot \mathbf{v}_R.$$

When the sign of the determinant changes, $\lambda_q$ is crossing the real axis, i.e., it is equal to zero and the above system of equations simplifies to

$$(\mathbf{J}_{DD} - q^2\mathbf{D}_{DD}) \cdot \mathbf{v}_D + (\mathbf{J}_{DR} - \mathbf{J}_{DI} \cdot \mathbf{J}_{II}^{-1} \cdot \mathbf{J}_{IR}) \cdot \mathbf{v}_R = \lambda_q \cdot \mathbf{v}_D = \mathbf{0},$$

$$\mathbf{J}_{RD} \cdot \mathbf{v}_D + (\mathbf{J}_{RR} - \mathbf{J}_{RI} \cdot \mathbf{J}_{II}^{-1} \cdot \mathbf{J}_{IR}) \cdot \mathbf{v}_R = \lambda_q \cdot \mathbf{v}_R = \mathbf{0},$$

which corresponds to the eigenvalue equation of the reduced model corresponding to the matrix

$$\begin{pmatrix} \mathbf{J}_{DD} - q^2\mathbf{D}_{DD} & \mathbf{J}_{DR} - \mathbf{J}_{DI}\mathbf{J}_{II}^{-1}\mathbf{J}_{IR} \\ \mathbf{J}_{RD} & \mathbf{J}_{RR} - \mathbf{J}_{RI}\mathbf{J}_{II}^{-1}\mathbf{J}_{IR} \end{pmatrix},$$

which you will notice takes exactly the same form as $\widetilde{\mathbf{M}}$. Thus, if $\begin{pmatrix}\mathbf{v}_D & \mathbf{v}_R & \mathbf{v}_I\end{pmatrix}^T$ is an eigenvector of the full model $\mathbf{M}$ corresponding to $\lambda_q$, then $\begin{pmatrix}\mathbf{v}_D & \mathbf{v}_R\end{pmatrix}^T$ is the eigenvector of the reduced model $\widetilde{\mathbf{M}}$ at the sign-change points. Because the distribution of phases is determined by the signs of the eigenvector elements, we deduce that the phases of all species in the reduced model are the same as in the full model. This holds at sign change points and, by continuity, sufficiently close to them. Whether this properly applies more generally for all values of $q$ is beyond the scope of this paper.

#### 2.3 Simplifying the calculation of the necessary condition $\mathcal{N}$ via the intracellular response function $F$

Inspecting $\widetilde{\mathbf{M}}$, we see that there are four blocks that must be computed:

- $(\mathbf{J}_{DD} - q^2\mathbf{D}_{DD})$: this represents the diffusion of and interactions between the extracellular species, which are exactly equivalent to the corresponding terms in the full matrix $\mathbf{M}$, and thus straightforward to compute.
- $\mathbf{J}_{RD}$: this represents how the extracellular species regulate transmembrane receptors in the system, leading to the formation of activated receptor complexes. As above, these elements are straightforward to compute since they are equivalent to the corresponding terms in the full matrix $\mathbf{M}$.
- $(\mathbf{J}_{DS} \stackrel{\text{def}}{=} \mathbf{J}_{DR} - \mathbf{J}_{DI}\mathbf{J}_{II}^{-1}\mathbf{J}_{IR})$: this represents how the intracellular signalling, using a single effective component $S$ per signalling pathway, regulates the extracellular species.
- $(\mathbf{J}_{SS} \stackrel{\text{def}}{=} \mathbf{J}_{RR} - \mathbf{J}_{RI}\mathbf{J}_{II}^{-1}\mathbf{J}_{IR})$: this represents the interactions involving the effective intracellular signalling components alone.

At first glance, the terms $\mathbf{J}_{DS}$ and $\mathbf{J}_{SS}$ appear highly non-trivial to compute and require computation of complex terms such as $\mathbf{J}_{II}^{-1}$. However, we can use Lemma 2.1 from [11] to show that $\mathbf{J}_{DS}$ and $\mathbf{J}_{SS}$ may be computed in a simpler manner, without a detailed knowledge of $\mathbf{J}_{II}$, provided one can estimate/measure the steady-state response function $F_{\text{output}}(\text{input})$. We briefly outline the arguments of Lemma 2.1 below to illustrate the general idea, and refer the reader to [11] for more details.

Consider the system 1, which has $n_D$ diffusible species, $n_R$ receptors, and $n_I$ intracellular species, yielding a total of $n_t = n_D + n_R + n_I$ species. WLOG we can order variables in such a way that there are first $n_r = n_D + n_R$ variables corresponding to the reduced model, followed by the $n_I = n_t - n_r$ intracellular species i.e., $\mathbf{c} \stackrel{\text{def}}{=} (c_1, c_2, \ldots, c_{n_r}, c_{n_r+1}, \ldots, c_{n_t})$.

If we assume all intracellular species are at steady state, then this allows us to define the quasi-steady state values of these species as a function of the other variables. Concretely, this yields the functions $\bar{c}_i(c_1, \dots, c_{n_r})$ for $i = n_r + 1, \dots, n$, which satisfy

$$0 = f_i(c_1, \dots, c_{n_r}, \bar{c}_{n_r+1}(c_1, \dots, c_{n_r}), \dots, \bar{c}_n(c_1, \dots, c_{n_r})).$$

If the quasi-steady state assumption holds, these functions allow us to remove the explicit dependence on the intracellular species from the PDEs and therefore rewrite them in terms of the reduced variables alone:

$$\frac{\partial c_i}{\partial t} = f_i(c_1, \dots, c_{n_r}, \bar{c}_{n_r+1}(c_1, \dots, c_{n_r}), \dots, \bar{c}_n(c_1, \dots, c_{n_r})) + D_i \nabla^2 c_i,$$

for $i = 1, \dots, n_r$.

By the multivariable chain rule, we can compute the Jacobian term of the corresponding linearized reaction-diffusion system:

$$\mathbf{J}_{quasi-steady-state} = \begin{pmatrix} \frac{\partial f_1}{\partial c_1} + \sum_{r=n_r+1}^{n} \frac{\partial f_1}{\partial c_r} \frac{\partial \bar{c_r}}{\partial c_1} & \cdots & \frac{\partial f_1}{\partial c_{n_r}} + \sum_{r=n_r+1}^{n} \frac{\partial f_1}{\partial c_r} \frac{\partial \bar{c_r}}{\partial c_{n_r}} \\ \vdots & \ddots & \vdots \\ \frac{\partial f_{n_r}}{\partial c_1} + \sum_{r=n_r+1}^{n} \frac{\partial f_{n_r}}{\partial c_r} \frac{\partial \bar{c_r}}{\partial c_1} & \cdots & \frac{\partial f_{n_r}}{\partial c_{n_r}} + \sum_{r=n_r+1}^{n} \frac{\partial f_{n_r}}{\partial c_r} \frac{\partial \bar{c_r}}{\partial c_{n_r}} \end{pmatrix}. \tag{5}$$

It can then be shown (see Lemma 2.1 in [11]), that this Jacobian can be related to the full Jacobian as follows:

$$\mathbf{J}_{quasi-steady-state} = \begin{pmatrix} \mathbf{J}_{DD} & \mathbf{J}_{DR} - \mathbf{J}_{DI}\mathbf{J}_{II}^{-1}\mathbf{J}_{IR} \\ \mathbf{J}_{RD} & \mathbf{J}_{RR} - \mathbf{J}_{RI}\mathbf{J}_{II}^{-1}\mathbf{J}_{IR} \end{pmatrix}$$

and hence that $\mathbf{J}_{quasi-steady-state}$ and $\widetilde{\mathbf{J}}$ are exactly equivalent. Therefore, $\widetilde{\mathbf{J}}$ (and hence $\widetilde{\mathbf{M}}$) need not be explicitly computed, but may be derived by assuming that all intracellular variables have reached a quasi-steady state, greatly simplifying the calculation of the terms $\mathbf{J}_{DS}$ and $\mathbf{J}_{SS}$. This means that the necessary condition $\boldsymbol{\mathcal{N}}$ may be computed without a detailed characterisation of intracellular dynamics.

##### 2.3.1 Biological interpretation of the terms $\mathbf{J}_{DS}$ and $\mathbf{J}_{SS}$

$\mathbf{J}_{DS}$ may be computed by characterising (e.g., via experiment) the steady-state (i.e., long-time limit) input-output relationship of the intracellular dynamics. Therefore, $\mathbf{J}_{DS}$ corresponds to the Jacobian terms of the associated intracellular response function $F_{\text{output}}(\text{input})$.

The term $\mathbf{J}_{SS}$ is less straightforward to interpret, and corresponds to the effective dynamics of the receptors/signalling when all intracellular species are assumed to be at steady state. There are three scenarios to consider:

- Case I: If the receptor species are not directly affected by intracellular elements, then $\mathbf{J}_{RI} = \mathbf{0}$, and thus $\mathbf{J}_{SS} = \mathbf{J}_{RR}$ corresponds to the degradation rates of the receptors and is independent of any intracellular parameters.
- Case II: $\mathbf{J}_{RI}$ is non-zero in such a way that the signs of corresponding elements in $\mathbf{J}_{SS}$ and $\mathbf{J}_{RR}$ are the same. In this case, the terms of $\mathbf{J}_{SS}$ correspond to the effective degradation rates of $S$, which may be impacted by both the degradation rates of the receptors and feedbacks from the intracellular species to the receptors.
- Case III: $\mathbf{J}_{RI}$ is nonzero in such a way that there exists at least a pair of corresponding elements in $\mathbf{J}_{SS}$ and $\mathbf{J}_{RR}$ with opposite signs. For example, a system with strong positive feedback between receptors and intracellular species could overcome the degradation rate of the receptor and lead to an effective auto-activation of species $S$ in the final reduced model.

Regardless of these scenarios, we have found (next chapter) that the necessary condition $\boldsymbol{\mathcal{N}}$ does not typically depend on the values of $\mathbf{J}_{SS}$ and therefore the biological interpretation of this term is often not required.

##### 2.4 Limitations of the necessary condition $\mathcal{N}$

The necessary condition $\mathcal{N}$ is necessary but not sufficient for Turing patterns to form. Here, we outline the scenarios in which the necessary condition $\mathcal{N}$ is not sufficient to guarantee symmetry breaking:

- **The system is unstable at long wavelengths ($q = 0$):** This would occur when $\mathbf{J}$ is unstable (i.e., at least one eigenvalue has positive real part), which cannot be deduced by considering $\widetilde{\mathbf{J}}$ alone.
- **The system is unstable at short wavelengths** ($q \to \infty$): This would occur if the following matrix were unstable: $\begin{pmatrix} \mathbf{J}_{RR} & \mathbf{J}_{RI} \\ \mathbf{J}_{IR} & \mathbf{J}_{II} \end{pmatrix}$ (see [11]), which also cannot be deduced by considering $\widetilde{\mathbf{J}}$ alone.
- **The system is unstable to oscillations:**

  Oscillations would be associated with a pair of complex conjugate eigenvalues crossing to the right half of the complex plane. In such a scenario, the sign of the signed determinant remains unchanged, and so cannot be detected by the necessary condition $\mathcal{N}$ alone. Of note, it has been shown that for models of size smaller or equal to 3, the necessary condition $\mathcal{N}$ is sufficient to guarantee that the instability that occurs is Turing [5]. Moreover, based on numerical analysis, it was suggested that this could hold for systems of size $\leq 5$ [5]. This agrees with our numerical results, as all the models of sizes 3, 4 and 5 we studied did not exhibit any other types of instabilities. However, a detailed analytical understanding of this problem requires further attention and is beyond the scope of this paper.

### 3 Analysis of activator-inhibitor models

#### 3.1 Case 1: The inhibitor binds to the ligand, inactivating it and targeting it for degradation

##### 3.1.1 Model formulation

In this scenario, the inhibitor irreversibly sequesters the activator by directly binding to it. Intracellular signalling and receptor species are represented by a single molecule $S$ (as justified in the previous section). We describe this behaviour in general terms by the following system of partial differential equations (PDEs):

$$\frac{\partial[A]}{\partial t} = F_A([S]) - k_+[A][I] - \delta_A[A] + D_A\nabla^2[A],$$
$$\frac{\partial[I]}{\partial t} = F_I([S]) - k_+[A][I] - \delta_I[I] + D_I\nabla^2[I],$$
$$\frac{\partial[S]}{\partial t} = F_S([A]) - \delta_S[S], \tag{6}$$

where $[\dots]$ denotes the concentration of each molecular species, which varies in space and time; $F$ stands for input-output overall intracellular response function; $k_+$ is the rate of formation of the complex; $\delta$ represents the degradation rate and $D$ stands for the diffusion rate. We assume that $F_S$ is linear.

##### 3.1.2 Linearizing the model about the steady state

To analyse the model, we first determine the steady state values $\bar{A}, \bar{I}, \bar{S}$ to find that:

$$F_A(\bar{S}) = k_+\bar{A}\bar{I} + \delta_A\bar{A},$$
$$F_I(\bar{S}) = k_+\bar{A}\bar{I} + \delta_I\bar{I},$$
$$F_S(\bar{A}) = \delta_S\bar{S},$$

We then linearize the system of PDEs close to equilibrium, i.e. where $[A] = \bar{A} + \Delta A$, $[I] = \bar{I} + \Delta I$ and $[S] = \bar{S} + \Delta S$. To normalize the deviation of concentrations about the steady state, we divide each of these equations by steady-state concentration to obtain

$$\frac{[A]}{\bar{A}} = 1 + \frac{\Delta A}{\bar{A}},\ \frac{[I]}{\bar{I}} = 1 + \frac{\Delta I}{\bar{I}} \text{ and } \frac{[S]}{\bar{S}} = 1 + \frac{\Delta S}{\bar{S}}.$$

Redefining $\Delta a \stackrel{\text{def}}{=\!=} \Delta A/\bar{A}$, $\Delta i \stackrel{\text{def}}{=\!=} \Delta I/\bar{I}$ and $\Delta s \stackrel{\text{def}}{=\!=} \Delta S/\bar{S}$ yields

$$\Delta a = \frac{[A]-\bar{A}}{\bar{A}},\ \Delta i = \frac{[I]-\bar{I}}{\bar{I}},\ \Delta s = \frac{[S]-\bar{S}}{\bar{S}}.$$

Substituting this into the system of equations 6, we obtain:

$$\frac{\partial}{\partial t}\Delta a = (\delta_A + k_+\bar{I})H_{S\to A}\Delta s - k_+\bar{I}(\Delta a + \Delta i) - \delta_A\Delta a + D_A\nabla^2(\Delta a),$$
$$\frac{\partial}{\partial t}\Delta i = (\delta_I + k_+\bar{A})H_{S\to I}\Delta s - k_+\bar{A}(\Delta a + \Delta i) - \delta_I\Delta i + D_I\nabla^2(\Delta i),$$
$$\frac{\partial}{\partial t}\Delta s = \delta_S\Delta a - \delta_S\Delta s, \tag{7}$$

where

$$H_{S\to A} \stackrel{\text{def}}{=\!=} \frac{\bar{S}}{F_A(\bar{S})}\frac{\partial F_A}{\partial S} \approx \frac{\Delta F_A}{F_A}\Big/\frac{\Delta S}{S} \tag{8}$$

is defined as the normalised sensitivity of the intracellular response function [13]. It expresses the (fractional) change in output for a given (fractional) change in input, see Figure 3A in main text. Intuitively, we may interpret $H_{S\to A} = 2$ to mean *'if the concentration of activated receptors increases by 1 percent, then the production rate of the receptor, $F_A(S)$, increases by 2 percent.* $H_{S\to I}$ is defined similarly. Notice here that when $F$ is the Hill function, $H$ is approximately equal to the Hill coefficient $n$:

$$H_{S\to A} = \frac{\bar{S}(K^n + \bar{S}^n)}{k\cdot \bar{S}^n} \cdot \frac{\partial}{\partial S}\left(\frac{k\cdot \bar{S}^n}{K^n+\bar{S}^n}\right) = n\cdot \frac{K^n}{K^n+\bar{S}^n} \approx n,$$

when $\bar{S}$ is small.

For ease of analysis, we then nondimensionalise 7 to obtain the following system

$$\begin{aligned} \frac{\partial}{\partial t}\Delta a &= H_{S\to A}\Delta s - k_{+I}(\Delta a + \Delta i - H_{S\to A}\Delta s) - \Delta a + \nabla^2(\Delta a), \\ \frac{\partial}{\partial t}\Delta i &= \delta_{I/A} H_{S\to I}\Delta s - k_{+A}(\Delta a + \Delta i - H_{S\to I}\Delta s) - \delta_{I/A}\Delta i + D_{I/A}\nabla^2(\Delta a), \\ \frac{\partial}{\partial t}\Delta s &= \delta_{S/A}\Delta a - \delta_{S/A}\Delta s, \end{aligned} \tag{9}$$

where

$$k_{+I} \overset{\text{def}}{=\!=} \frac{k_+\bar{I}}{\delta_A},\ k_{+A} \overset{\text{def}}{=\!=} \frac{k_+\bar{A}}{\delta_A},\ \delta_{I/A} \overset{\text{def}}{=\!=} \frac{\delta_I}{\delta_A},\ \delta_{S/A} \overset{\text{def}}{=\!=} \frac{\delta_S}{\delta_A},\ D_{I/A} \overset{\text{def}}{=\!=} \frac{D_I}{D_A}.$$

To examine the instability of the steady state to periodic disturbances, we apply a Fourier transform, yielding:

$$\frac{\partial}{\partial t}\begin{pmatrix}\Delta a_q\\ \Delta i_q\\ \Delta s_q\end{pmatrix} = \begin{pmatrix} -k_{+I}-1-q^2 & -k_{+I} & H_{S\to A}(1+k_{+I}) \\ -k_{+A} & -k_{+A}-\delta_{I/A}-D_{I/A}q^2 & H_{S\to I}(\delta_{I/A}+k_{+A}) \\ \delta_{S/A} & 0 & -\delta_{S/A}\end{pmatrix}\begin{pmatrix}\Delta a_q\\ \Delta i_q\\ \Delta s_q\end{pmatrix}. \tag{10}$$

Hence, the reaction-diffusion matrix we need to test for the necessary condition $\boldsymbol{\mathcal{N}}$ takes the form

$$\widetilde{\mathbf{M}}(q^2) = \begin{pmatrix} -k_{+I}-1-q^2 & -k_{+I} & H_{S\to A}(1+k_{+I}) \\ -k_{+A} & -k_{+A}-\delta_{I/A}-D_{I/A}q^2 & H_{S\to I}(\delta_{I/A}+k_{+A}) \\ \delta_{S/A} & 0 & -\delta_{S/A}\end{pmatrix}.$$

##### 3.1.3 Deriving the necessary condition $\boldsymbol{\mathcal{N}}$ for pattern formation

The necessary condition $\boldsymbol{\mathcal{N}}$ states that the determinant of the linearized reaction-diffusion matrix must change sign twice - at $q_0$ and $q_1$, two real wavenumbers. We find (see below) that this determinant will take the following polynomial form: $a_2q^4 + a_1q^2 + a_0$, with $a_2 > 0$. The condition $\boldsymbol{\mathcal{N}}$ implies that $a_0 > 0$ and $a_1 < 0$ to ensure the system is capable of changing sign twice; and $a_1^2 - 4a_2a_0 > 0$ to ensure the sign changes at real (and not complex) values of $q$. These restrictions on the coefficients of the polynomial may be derived from Descartes' law of signs (see, for example, [14], Chapter IV, or [5]).

For this particular model we evaluate

$$\begin{aligned}(-1)^n det(\widetilde{\mathbf{M}}(q^2)) &= D_{I/A}\delta_{S/A}\cdot q^4 - \left(\delta_{S/A}\left(D_{I/A}(H_{S\to A}-1)(1+k_{+I}) - \delta_{I/A} - k_{+A}\right)\right)\cdot q^2 + \\ &- \delta_{S/A}\left((\delta_{I/A}+k_{+A}+\delta_{I/A}k_{+I})(H_{S\to A}-1) + k_{+I}\left(H_{S\to A}k_{+A} - H_{S\to I}(\delta_{I/A}+k_{+A})\right)\right). \end{aligned} \tag{11}$$

Notice that coefficient corresponding to $q^4$ is always non-negative: $D_{I/A}\delta_{S/A} \geq 0$.
To satisfy the necessary condition $\boldsymbol{\mathcal{N}}$, expression 11 must be positive at $q = 0$, i.e.,

$$-\delta_{S/A}\left((\delta_{I/A}+k_{+A}+\delta_{I/A}k_{+I})(H_{S\to A}-1) + k_{+I}\left(H_{S\to A}k_{+A} - H_{S\to I}(\delta_{I/A}+k_{+A})\right)\right) > 0. \tag{12}$$

For 11 to turn negative, the coefficient corresponding to $q^2$ must be negative and therefore

$$\left(\delta_{S/A}\left(D_{I/A}(H_{S\to A}-1)(1+k_{+I}) - \delta_{I/A} - k_{+A}\right)\right) > 0. \tag{13}$$

Inequality 13 implies that $H_{S\to A} > 1$, in particular

$$H_{S\to A} - 1 > \frac{\delta_{I/A} + k_{+A}}{D_{I/A}(1 + k_{+I})} > 0. \tag{14}$$

To satisfy inequality 12, we need

$$(\delta_{I/A} + k_{+A} + \delta_{I/A}k_{+I})(H_{S\to A} - 1) < -k_{+I}\left(H_{S\to A}k_{+A} - H_{S\to I}(\delta_{I/A} + k_{+A})\right),$$

which is equivalent to

$$(k_{+I}k_{+A} + \delta_{I/A} + k_{+A} + \delta_{I/A}k_{+I})(H_{S\to A}) - (\delta_{I/A} + k_{+A} + \delta_{I/A}k_{+I}) < H_{S\to I}k_{+I}(\delta_{I/A} + k_{+A}),$$

we can expand this expression to

$$\frac{(k_{+I}k_{+A} + \delta_{I/A}k_{+I})(H_{S\to A})}{k_{+I}(\delta_{I/A} + k_{+A})} + \frac{(\delta_{I/A} + k_{+A})(H_{S\to A})}{k_{+I}(\delta_{I/A} + k_{+A})} - \frac{(\delta_{I/A} + k_{+A} + \delta_{I/A}k_{+I})}{k_{+I}(\delta_{I/A} + k_{+A})} < H_{S\to I},$$

which is equivalent to

$$H_{S\to A} + \frac{(\delta_{I/A} + k_{+A})(H_{S\to A})}{k_{+I}(\delta_{I/A} + k_{+A})} - \frac{\delta_{I/A} + k_{+A}}{k_{+I}(\delta_{I/A} + k_{+A})} - \frac{\delta_{I/A}k_{+I}}{k_{+I}(\delta_{I/A} + k_{+A})} < H_{S\to I},$$

and, using $H_{S\to A} - 1 > 0$, we conclude that

$$H_{S\to A} - 1 < H_{S\to A} + (H_{S\to A} - 1)\cdot\frac{1}{k_{+I}} - \frac{\delta_{I/A}}{\delta_{I/A} + k_{+A}} < H_{S\to I} \tag{15}$$

as, clearly,

$$-1 < (H_{S\to A} - 1)\cdot\frac{1}{k_{+I}} - \frac{\delta_{I/A}}{\delta_{I/A} + k_{+A}}.$$

Therefore, regardless of parameters, this model may satisfy the necessary condition $\boldsymbol{\mathcal{N}}$ only if $H_{S\to A} - 1 > 0$ and $H_{S\to A} - 1 < H_{S\to I}$. These are the general limits that are plotted in the main text (light blue lines in Figures 3,4).

Notice that we did not write out in detail the condition $a_1^2 - 4a_2a_0 > 0$, which must be satisfied for the roots $q_0, q_1$ to be real. However, when directly comparing the necessary condition $\boldsymbol{\mathcal{N}}$ to the necessary and sufficient conditions for patterning, we compute this additional inequality for $\boldsymbol{\mathcal{N}}$ numerically.

#### 3.2 Case 2: The inhibitor binds to the receptor, competing with the ligand for pathway activation

##### 3.2.1 Model formulation

In this scenario, the inhibitor competes with the activator for binding to the receptor. Intracellular signalling and receptor species are again represented by a single molecule $S$. We describe this behaviour in general terms by the following system of partial differential equations (PDEs):

$$\begin{aligned}
\frac{\partial[A]}{\partial t} &= F_A([S]) - \delta_A[A] + D_A\nabla^2[A],\\
\frac{\partial[I]}{\partial t} &= F_I([S]) - \delta_I[I] + D_I\nabla^2[I],\\
\frac{\partial[S]}{\partial t} &= F_S([A],[I]) - \delta_S[S],
\end{aligned} \tag{16}$$

where $[\dots]$ denotes concentration of each molecular species, which varies in space and time; $F$ stands for the input-output overall intracellular response function; $\delta$ stands for the degradation rate and $D$ stands for the diffusion rate. We assume that $F_S \propto [A]/[I]$.

##### 3.2.2 Linearizing the model about the steady state

As before, we find the steady states $\bar{A}, \bar{I}, \bar{S}$ via:

$$F_A(\bar{S}) = \delta_A \bar{A},$$
$$F_I(\bar{S}) = \delta_I \bar{I},$$
$$F_S(\bar{A}, \bar{I}) = \delta_S \bar{S},$$

We then linearise the system of PDEs close to equilibrium and define new variables $\Delta a$, $\Delta i$, $\Delta s$, which describe the normalised deviation of concentrations about the steady state, i.e., $\Delta a = ([A] - \bar{A})/\bar{A}$.

Substituting this into the system of equations 14, we obtain:

$$\begin{aligned} \frac{\partial}{\partial t}\Delta a &= \delta_A H_{S\to A}\Delta - \delta_A \Delta a + D_A \nabla^2(\Delta a), \\ \frac{\partial}{\partial t}\Delta i &= \delta_I H_{S\to I}\Delta s - \delta_I \Delta i + D_I \nabla^2(\Delta i), \\ \frac{\partial}{\partial t}\Delta s &= \delta_S \Delta a - \delta_S \Delta i - \delta_S \Delta s, \end{aligned} \tag{17}$$

where $H_{S\to A}$ and $H_{S\to A}$ are defined as in the previous section (see equation 8).

We then nondimensionalise to obtain the following system:

$$\begin{aligned} \frac{\partial}{\partial t}\Delta a &= H_{S\to A}\Delta s - \Delta a + \nabla^2(\Delta a), \\ \frac{\partial}{\partial t}\Delta i &= \delta_{I/A} H_{S\to I}\Delta s - \delta_{I/A} \Delta i + D_{I/A} \nabla^2(\Delta a), \\ \frac{\partial}{\partial t}\Delta s &= \delta_{S/A} \Delta a - \delta_{S/A} \Delta i - \delta_{S/A} \Delta s, \end{aligned} \tag{18}$$

where

$$\delta_{I/A} \stackrel{\text{def}}{=\!=} \frac{\delta_I}{\delta_A}, \ \delta_{S/A} \stackrel{\text{def}}{=\!=} \frac{\delta_S}{\delta_A}, \text{ and } D_{I/A} \stackrel{\text{def}}{=\!=} \frac{D_I}{D_A}.$$

This gives the following reaction-diffusion matrix to test for the necessary condition $\boldsymbol{\mathcal{N}}$:

$$\widetilde{\mathbf{M}}(q^2) = \begin{pmatrix} -1 - q^2 & 0 & H_{S\to A} \\ 0 & -\delta_{I/A} - D_{I/A} \cdot q^2 & H_{S\to I}\delta_{I/A} \\ \delta_{S/A} & -\delta_{S/A} & -\delta_{S/A} \end{pmatrix}.$$

##### 3.2.3 Deriving necessary condition $\boldsymbol{\mathcal{N}}$ for pattern formation

We evaluate

$$\begin{aligned} (-1)^n det(\widetilde{\mathbf{M}}(q^2)) = D_{I/A} \cdot \delta_{S/A} \cdot q^4 + \left(\delta_{S/A}\left(D_{I/A}(1 - H_{S\to A}) + \delta_{I/A}(1 + H_{S\to I})\right)\right) \cdot q^2 + \\ + \delta_{S/A}\delta_{I/A}\left(1 - H_{S\to A} + H_{I\to S}\right). \end{aligned} \tag{19}$$

As before, this is a polynomial of the form $a_2 q^4 + a_1 q^2 + a_0$, with $a_2 > 0$, whose coefficients must satisfy $a_0 > 0$, $a_1 < 0$ to change sign twice for positive $q$.

To satisfy the necessary condition $\boldsymbol{\mathcal{N}}$, expression 19 must be positive at $q = 0$, i.e.,

$$H_{S\to A} - 1 < H_{S\to I}. \tag{20}$$

For 19 to turn negative, by Descartes' rule of signs, the coefficient corresponding to $q^2$ must be negative and therefore

$$H_{S\to A} - 1 > \frac{\delta_{I/A}}{D_{I/A}}(1 + H_{S\to I}) \tag{21}$$

implying:

$$\frac{H_{S\to A}-1}{1+H_{S\to I}}>0 \tag{22}$$

We can see that inequalities 20 and 22 may be simultaneously satisfied in two cases:

- Case 1: $H_{S\to A}>1$ and $H_{S\to I}>0$ (together with inequality 20)
- Case 2: $H_{S\to A}<0$ and $H_{S\to I}<-1$ (together with inequality 20)

These inequalities are general limits that apply regardless of parameters and are plotted as light blue lines in the main text figures. As before, we compute an additional inequality ($a_1^2-4a_2a_0>0$) numerically when explicitly comparing the necessary condition $\boldsymbol{\mathcal{N}}$ to the necessary and sufficient conditions.

###### 3.2.4 Explicitly considering binding of inhibitor to receptor

Strictly speaking, to precisely model the scenario in which the inhibitor binds to the receptor and competes with the ligand for pathway activation, we should explicitly consider the (immobile) complex formed between the inhibitor and the receptor, rather than modelling signalling as being directly inhibited by the inhibitor, as we have implicitly assumed in the previous section. Here we shall show that the restrictions on the regulatory feedbacks in this type of model are qualitatively similar whether or not the inhibitor-receptor binding is explicitly accounted for.

To do this, we consider the dynamics of the the receptor $R$, the receptor-inhibitor complex $C$, and the activated receptor complex (i.e., signalling) $S$, via the following system of partial differential equations (PDEs):

$$\begin{aligned}
\frac{\partial[A]}{\partial t}&=F_A([S])-\delta_A[A]+D_A\nabla^2[A],\\
\frac{\partial[I]}{\partial t}&=F_I([S])-\delta_I[I]+D_I\nabla^2[I]-k_+[I][R]+k_-[C],\\
\frac{\partial[S]}{\partial t}&=\lambda_S[A][R]-\delta_S[S],\\
\frac{\partial[R]}{\partial t}&=\lambda_R-\delta_R[R]-k_+[I][R]+k_-[C],\\
\frac{\partial[C]}{\partial t}&=k_+[I][R]-k_-[C]-\delta_C[C].
\end{aligned} \tag{23}$$

Through the same steps as before, we derive the linearized reaction-diffusion matrix

$$\mathbf{M}(q^2)=\begin{pmatrix}
-\delta_A-D_Aq^2 & 0 & \delta_A H_{S\to A} & 0 & 0\\
0 & -\delta_I-D_I\cdot q^2-k_+\bar{R} & (\delta_I+\delta_C\frac{k_+\bar{R}}{\delta_C+k_-})H_{S\to I} & -k_+\bar{R} & k_-\frac{k_+\bar{R}}{\delta_C+k_-}\\
\delta_S & 0 & -\delta_S & \delta_S & 0\\
0 & -k_+\bar{I} & 0 & -k_+\bar{I}-\delta_R & k_-\frac{k_+\bar{I}}{\delta_C+k_-}\\
0 & k_-+\delta_C & 0 & k_-+\delta_C & -k_--\delta_C
\end{pmatrix}.$$

Since $C$ and $R$ do not diffuse between cells, we can use the Schur complement approach once more to simplify the calculation of $\boldsymbol{\mathcal{N}}$. In particular, we may compute $\boldsymbol{\mathcal{N}}$ from a reduced model:

$$\widetilde{\mathbf{M}}=\begin{pmatrix}
-D_Aq^2-\delta_A & 0 & H_{S\to A}\delta_A\\
0 & -\bar{\delta_I}-D_I\cdot q^2 & H_{S\to I}\bar{\delta_I}\left(1+\frac{K_1}{\bar{\delta_I}}\cdot K_2\right)\\
\delta_S & -\delta_S K_2 & -\delta_S
\end{pmatrix}$$

where

$$\bar{\delta_I} \stackrel{\text{def}}{=} \delta_I + \frac{\bar{R}\delta_C\delta_R k_+}{\delta_C\delta_R + \delta_R k_- + \bar{I}\delta_C k_+},$$

$$K_1 \stackrel{\text{def}}{=} \frac{\bar{R}k_+}{1 + k_-/\delta_C} = \frac{\bar{R}k_+\delta_R\delta_C}{\delta_C\delta_R + k_-\delta_R},$$

$$K_2 \stackrel{\text{def}}{=} \frac{\bar{I}\delta_C k_+}{\delta_C\delta_R + \delta_R k_- + \bar{I}\delta_C k_+} \leq 1.$$

Redefining: $C_1 \stackrel{\text{def}}{=} 1 + \frac{K_1}{\bar{\delta_I}} \cdot K_2 > 1$ and non-dimensionalizing, we find:

$$\widetilde{\mathbf{M}} = \begin{pmatrix} -q^2 - 1 & 0 & H_{S\to A} \\ 0 & -\overline{\delta_{I/A}} - D_{I/A} \cdot q^2 & H_{S\to I}\overline{\delta_{I/A}}C_1 \\ \delta_{S/A} & -\delta_{S/A}K_2 & -\delta_{S/A} \end{pmatrix}.$$

Computing the determinant gives the following:

$$(-1)^n det(\widetilde{\mathbf{M}}(q^2)) = D_{I/A} \cdot \delta_{S/A} \cdot q^4 + \left(\delta_{S/A}\left(D_{I/A}(1 - H_{S\to A}) + \delta_{I/A}(1 + H_{I\to S}C_1K_2)\right)\right) \cdot q^2 + \\ + \delta_{S/A}\delta_{I/A}\left(1 - H_{S\to A} + H_{I\to S}C_1K_2\right). \tag{24}$$

To satisfy the necessary condition, the following two inequalities must hold:

$$H_{S\to A} - 1 < H_{S\to I}C_1K_2. \tag{25}$$

$$\frac{H_{S\to A} - 1}{1 + H_{S\to I}C_1K_2} > 0. \tag{26}$$

These take a similar form to the simpler model, but with $H_{S\to I}$ rescaled by a factor of $C_1K_2$. There are again two cases:

- Case 1: $H_{S\to A} > 1$ and $H_{S\to I} > 0$ (together with inequality 25)
- Case 2: $H_{S\to A} < 0$ and $H_{S\to I}C_1K_2 < -1$ (together with inequality 25)

We can see that the same qualitative (i.e., sign) restrictions on $H_{S\to A}$ and $H_{S\to I}$ must still hold: either both are positive or both are negative.

Whilst of the same form, the *quantitative* values of $H_{S\to I}$ restricted by the necessary condition $\boldsymbol{\mathcal{N}}$ are different. In the limit when $K_2 \approx 1$ (i.e., when the inhibitor is in excess: $(k_- + \delta_C)\delta_R << \delta_C k_+ \bar{I}$) and $K_1/\bar{\delta_1} \ll 1$, the $H_{S\to A}$-$H_{S\to I}$ region restricted by $\boldsymbol{\mathcal{N}}$ will be very similar to the one for simpler model.

#### 3.3 Case 3: The inhibitor reversibly binds the ligand to form a complex which cannot activate signalling but can dissociate to release the ligand

##### 3.3.1 Model formulation

In this scenario, the inhibitor reversibly sequesters the activator by binding directly to it and forming a complex $C$. Intracellular signalling and receptor species are represented by a single molecule $S$, leading to the following system of PDEs:

$$\frac{\partial[A]}{\partial t} = F_A([S]) + k_-[C] - k_+[A][I] - \delta_A[A] + D_A\nabla^2[A],$$

$$\frac{\partial[I]}{\partial t} = F_I([S]) + k_-[C] - k_+[A][I] - \delta_I[I] + D_I\nabla^2[I],$$

$$\frac{\partial[C]}{\partial t} = -k_-[C] + k_+[A][I] + D_C\nabla^2[C],$$

$$\frac{\partial[S]}{\partial t} = F_S([A]) - \delta_S[S], \tag{27}$$

where $[\dots]$ denotes the concentration of each molecular species, which varies in space and time; $F$ stands for the input-output overall intracellular response function; $k_+$ is the rate of formation of the complex; $k_-$ is the rate of dissociation of the complex; $\delta$ represents the rate of degradation and $D$ represents the rate of diffusion. We assume that $F_S$ is linear.

##### 3.3.2 Linearizing the model about the steady state

The steady states $\bar{A}, \bar{I}, \bar{C}, \bar{S}$ satisfy:

$$F_A(\bar{S}) = \delta_A\bar{A},$$

$$F_I(\bar{S}) = \delta_I\bar{I},$$

$$k_-\bar{C} = k_+\bar{A}\bar{I}$$

$$F_S(\bar{A}) = \delta_S\bar{S},$$

Close to the steady state, we obtain:

$$\frac{\partial}{\partial t}\Delta a = \delta_A H_{S\to A}\Delta s - \delta_A\Delta a + k_+\bar{I}(\Delta c - \Delta i - \Delta a) + D_A\nabla^2(\Delta a),$$

$$\frac{\partial}{\partial t}\Delta i = \delta_I H_{S\to I}\Delta s - \delta_I\Delta i + k_+\bar{A}(\Delta c - \Delta i - \Delta a) + D_I\nabla^2(\Delta i),$$

$$\frac{\partial}{\partial t}\Delta c = -k_-(\Delta c - \Delta i - \Delta a) + D_C\nabla^2(\Delta c),$$

$$\frac{\partial}{\partial t}\Delta s = \delta_S\Delta a - \delta_S\Delta s, \tag{28}$$

where $H_{S\to A}$ and $H_{S\to A}$ are defined as in the previous section (see equation 8).

Following non-dimensionalization, we obtain the following system:

$$\frac{\partial}{\partial t}\Delta a = H_{S\to A}\Delta s - \Delta a + k_{+I}(\Delta c - \Delta i - \Delta a) + \nabla^2(\Delta a),$$

$$\frac{\partial}{\partial t}\Delta i = \delta_{I/A}H_{S\to I}\Delta s - \delta_{I/A}\Delta i + k_{+A}(\Delta c - \Delta i - \Delta a) + D_{I/A}\nabla^2(\Delta i),$$

$$\frac{\partial}{\partial t}\Delta c = -k_{-A}(\Delta c - \Delta i - \Delta a) + D_{C/A}\nabla^2(\Delta c),$$

$$\frac{\partial}{\partial t}\Delta s = \delta_{S/A}\Delta a - \delta_{S/A}\Delta s, \tag{29}$$

where

$$k_{+I} \overset{\text{def}}{=\!=} \frac{k_+\bar{I}}{\delta_A},\ k_{+A} \overset{\text{def}}{=\!=} \frac{k_+\bar{A}}{\delta_A},\ k_{-A} \overset{\text{def}}{=\!=} \frac{k_-}{\delta_A},\ \delta_{I/A} \overset{\text{def}}{=\!=} \frac{\delta_I}{\delta_A},\ \delta_{S/A} \overset{\text{def}}{=\!=} \frac{\delta_S}{\delta_A},\ D_{I/A} \overset{\text{def}}{=\!=} \frac{D_I}{D_A},\ D_{C/A} \overset{\text{def}}{=\!=} \frac{D_C}{D_A}.$$

This yields the following reaction-diffusion matrix to test for the necessary condition $\boldsymbol{\mathcal{N}}$:

$$\widetilde{\mathbf{M}}(q^2) = \begin{pmatrix} -1 - k_{+I} - q^2 & -k_{+I} & k_{+I} & H_{S\to A} \\ -k_{+A} & -\delta_{I/A} - k_{+A} - D_{I/A}\cdot q^2 & k_{+A} & H_{S\to I}\delta_{I/A} \\ k_{-A} & k_{-A} & -k_{-A} - D_{C/A}\cdot q^2 & 0 \\ \delta_{S/A} & 0 & 0 & -\delta_{S/A} \end{pmatrix}.$$

##### 3.3.3 Deriving the necessary condition $\mathcal{N}$ for pattern formation

In this case, the determinant of $\widetilde{\mathbf{M}}(q^2)$ takes the form $a_3q^6+a_2q^4+a_1q^2+a_0$, with $a_3 > 0$. In order to change sign twice, Descartes' law of signs requires $a_0 > 0$ and at least one of $a_2$, $a_1$ to be negative. Further, the discriminant must be positive: $a_2^2a_1^2 - 4a_3a_1^3 - 4a_2^3a_0 - 27a_3^2a_0^2 + 18a_3a_2a_1a_0 > 0$ to ensure the sign changes at real (and not complex) values of $q^2$.

For this particular model we evaluate

$$\begin{aligned}
(-1)^n det(\widetilde{\mathbf{M}}(q^2)) = D_{I/A}D_{C/A}\delta_{S/A} \cdot q^6 + \\
+ \left(\delta_{S/A}\left(D_{I/A}D_{C/A}(1-H_{S\to A}) + D_{I/A}k_{-A} + D_{C/A}(\delta_{I/A} + k_{+A} + D_{I/A}k_{+I})\right)\right) \cdot q^4 \\
+ \left(\delta_{S/A}\left((1-H_{S\to A})(D_{C/A}\delta_{I/A} + D_{C/A}k_{+A} + D_{I/A}k_{-A}) + \delta_{I/A}k_{-A} + D_{C/A}\delta_{I/A}k_{+I}(1+H_{S\to I})\right)\right) \cdot q^2 \\
+ \delta_{S/A}\delta_{I/A}k_{-A}\left(1-H_{S\to A}\right).
\end{aligned} \tag{30}$$

To satisfy the necessary condition $\mathcal{N}$, expression 30 must be positive at $q = 0$, i.e.,

$$H_{S\to A} < 1. \tag{31}$$

This directly implies that the coefficient corresponding to $q^4$ is always positive (i.e., $H_{S\to I} < 1 \implies a_2 > 0$). Thus, to satisfy $\mathcal{N}$, the coefficient corresponding to $q^2$ must be negative ($a_1 < 0$). Therefore,

$$H_{S\to I} < -1. \tag{32}$$

Together, these inequalities ($H_{S\to A} < 1$ and $H_{S\to I} < -1$) are general limits that apply regardless of parameters and are plotted in light blue lines in the figures of the main text.

When comparing the necessary condition $\mathcal{N}$ to the necessary and sufficient conditions, we again compute these inequalities numerically, also incorporating the cubic discriminant into our calculations.

##### 3.3.4 The scenario in which the complex both degrades and dissociates

In general, the complex formed by the inhibitor and ligand could both degrade and dissociate at non-zero rates. For simplicity, we have so far assumed the limiting cases in which either degradation or dissociation is considered negligible. Here, we will show that it is the relative ratio between these two rates that determines the constraints on the regulatory functions. In particular, if degradation of the complex is much faster than its dissociation, then the results from Figure 4A will apply; if complex dissociation is much faster than its degradation, then the results from Figure 4C will apply; and for intermediate values of the dissociation:degradation ratio, regulatory circuits are possible in the space defined by the combination of Figure 4A and Figure 4C.

To explore this more general case, we consider the following set of equations:

$$\frac{\partial[A]}{\partial t} = F_A([S]) + k_-[C] - k_+[A][I] - \delta_A[A] + D_A\nabla^2[A],$$

$$\frac{\partial[I]}{\partial t} = F_I([S]) + k_-[C] - k_+[A][I] - \delta_I[I] + D_I\nabla^2[I],$$

$$\frac{\partial[C]}{\partial t} = -k_-[C] + k_+[A][I] - \delta_C[C] + D_C\nabla^2[C],$$

$$\frac{\partial[S]}{\partial t} = F_S([A]) - \delta_S[S].$$

We denote steady states as $\bar{A}$,$\bar{I}$,$\bar{C}$,$\bar{S}$, leading to:

$$\frac{F_A(\bar{S})}{\bar{A}} = \delta_A + k_+\bar{I}\frac{\delta_C}{\delta_C + k_-}$$

and

$$\frac{F_I(\bar{S})}{\bar{I}} = \delta_I + k_+\bar{A}\frac{\delta_C}{\delta_C + k_-}.$$

After linearizing, we obtain:

$$\frac{\partial}{\partial t}\Delta a = (\delta_A + k_+\bar{I}\frac{\delta_C}{\delta_C + k_-})H_{S\to A}\Delta s - \delta_A\Delta a + k_+\bar{I}(\frac{k_-}{k_- + \delta_C}\Delta c - \Delta i - \Delta a) + D_A\nabla^2(\Delta a),$$

$$\frac{\partial}{\partial t}\Delta i = (\delta_I + k_+\bar{A}\frac{\delta_C}{\delta_C + k_-})H_{S\to I}\Delta s - \delta_I\Delta i + k_+\bar{A}(\frac{k_-}{k_- + \delta_C}\Delta c - \Delta i - \Delta a) + D_I\nabla^2(\Delta i),$$

$$\frac{\partial}{\partial t}\Delta c = (-k_- - \delta_C)(\Delta c - \Delta i - \Delta a) + D_C\nabla^2(\Delta c),$$

$$\frac{\partial}{\partial t}\Delta s = \delta_S\Delta a - \delta_S\Delta s,$$

where $H_{S\to A}$ and $H_{S\to A}$ are defined as usual.

By inspection, we can see that the structure of these equations depends on the terms $\frac{\delta_C}{\delta_C+k_-}$ and $\frac{k_-}{k_-+\delta_C}$, which both depend on the ratio of degradation to dissociation rates, $\delta_C/k_-$. Therefore, when $\delta_C/k_- >> 1$, the equations have the same form as system 7 and thus the allowed patterning circuits correspond to Figure 4A. In contrast, when $\delta_C/k_- << 1$, the equations have the same form as system 28 and now the allowed circuits correspond to Figure 4C. We considered values of $\delta_C/k_-$ outside of the limiting cases numerically, and confirm our intuition that in general, circuits corresponding to both limiting cases can form patterns for intermediate values of $\delta_C/k_-$ (Figure S4). The implication of these results is that the degradation rate of the complex need not be identically zero for the non-intuitive circuits in Figure 4C to be applicable; it instead suffices that the complex dissociation rate is merely sufficiently high compared to its degradation rate.

We expect the value of $\delta_C/k_-$ to depend on the specific activator-inhibitor pair as well as the *in vivo* context. For complexes with very high affinity (equivalently, low values for the dissociation constant $K_d \equiv k_-/k_+$), it is likely that degradation would dominate over dissociation. However, there is good evidence to suggest that this would not be the case for all complexes, and there may be many which instead dissociate appreciably before being degraded (i.e., $\delta_C/k_-$ is small). Firstly, surface plasmon resonance (SPR) measurements have estimated complex dissociation rates *in vitro* and these can be relatively rapid (e.g., $1/k_-$ is on the order of minutes for BMP2-Chordin [15], BMP2/4-Follistatin [16], and WNT3a-sFRP1/2 [17]). Molecular degradation rates are tissue-dependent, but half-lives are often on the order of hours (e.g., [18, 19]). Therefore, it is feasible to expect that complex dissociation will be relevant for many of these cases. Moreover, whilst these dissociation rates are difficult to directly measure *in vivo*, there is indirect evidence of complex dissociation *in vivo* from the phenomenon of ligand shuttling observed in several systems. For example, the addition of sFRPs (WNT inhibitors) in Xenopus embryos leads to an increase in WNT signalling at long distances from the WNT source, which is only possible if the WNT-sFRP complex can readily dissociate [20] [21]. Taken together, these results indicate that complex dissociation need not be negligible compared to its degradation rate, and therefore that the circuits identified in Figure 4C are likely to be broadly applicable.

#### 3.4 Case 4: The inhibitor acts indirectly via a separate signalling pathway

##### 3.4.1 Model formulation

In this scenario, the inhibitor acts indirectly - there are two diffusible ligands, each activating a different signalling pathway, with transcriptional/intracellular feedbacks possible between the two pathways. The intracellular signalling and receptor species of each signalling pathway are represented by a single molecule $S$, i.e., $S_{A1}, S_{A2}$. We assume that the signalling pathways regulate each other and that one of them (WLOG we have chosen pathway 1) can also self-regulate itself. We describe this behaviour in general terms by the following system of PDEs:

$$
\begin{aligned}
\frac{\partial[A1]}{\partial t} &= F_{A1}([S_{A1}],[S_{A2}]) - \delta_{A1}[A1] + D_{A1}\nabla^2[A1],\\
\frac{\partial[A2]}{\partial t} &= F_{A2}([S_{A1}]) - \delta_{A2}[A2] + D_{A2}\nabla^2[A2],\\
\frac{\partial[S_{A1}]}{\partial t} &= F_{S_{A1}}([A1]) - \delta_{S_{A1}}[S_{A1}],\\
\frac{\partial[S_{A2}]}{\partial t} &= F_{S_{A2}}([A2]) - \delta_{S_{A2}}[S_{A2}],
\end{aligned} \tag{33}
$$

where $[\ldots]$ denotes the concentration of each molecular species, which varies in space and time; $F$ stands for input-output overall intracellular response function; $\delta$ stands for degradation rate and $D$ the diffusion rate. We assume that $F_{S_{A1}}, F_{S_{A2}}$ are both linear.

##### 3.4.2 Linearizing the model about the steady state

The steady states $\overline{A1}, \overline{A2}, \overline{S_{A1}}, \overline{S_{A2}}$ are given by:

$$
\begin{aligned}
F_{A1}(\overline{S_{A1}}, \overline{S_{A2}}) &= \delta_{A1}\overline{A1},\\
F_{A2}(\overline{S_{A1}}) &= \delta_{A2}\overline{A2},\\
F_{S_{A1}}(\overline{A1}) &= \delta_{S_{A1}}\overline{S_{A1}},\\
F_{S_{A2}}(\overline{A2}) &= \delta_{S_{A2}}\overline{S_{A2}},
\end{aligned}
$$

This yields the following equations for the linearized dynamics:

$$
\begin{aligned}
\frac{\partial}{\partial t}\Delta a_1 &= \delta_{A1}H_{S_{A1}\to A1}\Delta s_1 + \delta_{A1}H_{S_{A2}\to A1}\Delta s_2 - \delta_{A1}\Delta a_1 + D_{A1}\nabla^2(\Delta a_1),\\
\frac{\partial}{\partial t}\Delta a_2 &= \delta_{A2}H_{S_{A1}\to A2}\Delta s_1 - \delta_{A2}\Delta a_2 + D_{A2}\nabla^2(\Delta a_2),\\
\frac{\partial}{\partial t}\Delta s_1 &= \delta_{S_{A1}}\Delta a_1 - \delta_{S_{A1}}\Delta s_1,\\
\frac{\partial}{\partial t}\Delta s_2 &= \delta_{S_{A2}}\Delta a_2 - \delta_{S_{A2}}\Delta s_2,
\end{aligned} \tag{34}
$$

where $H_{S_{A1}\to A2}$ and $H_{S_{A2}\to A1}$ are defined as before (see equation 8).

Non-dimensionalization yields:

$$
\begin{aligned}
\frac{\partial}{\partial t}\Delta a_1 &= H_{S_{A1}\to A1}\Delta s_1 + H_{S_{A2}\to A1}\Delta s_2 - \Delta a_1 + \nabla^2(\Delta a_1),\\
\frac{\partial}{\partial t}\Delta a_2 &= \delta_{A2/A1}H_{S_{A1}\to A2}\Delta s_1 - \delta_{A2/A1}\Delta a_2 + D_{A2/A1}\nabla^2(\Delta a_2),\\
\frac{\partial}{\partial t}\Delta s_1 &= \delta_{S_{A1}/A1}\Delta a_1 - \delta_{S_{A1}/A1}\Delta s_1,\\
\frac{\partial}{\partial t}\Delta s_2 &= \delta_{S_{A2}/A1}\Delta a_2 - \delta_{S_{A2}/A1}\Delta s_2,
\end{aligned} \tag{35}
$$

where

$$
\delta_{A2/A1} \stackrel{\text{def}}{=} \frac{\delta_{A2}}{\delta_{A1}},\ \delta_{S_{A1}/A1} \stackrel{\text{def}}{=} \frac{\delta_{S_{A1}}}{\delta_{A1}},\ \delta_{S_{A2}/A1} \stackrel{\text{def}}{=} \frac{\delta_{S_{A2}}}{\delta_{A1}},\ D_{A2/A1} \stackrel{\text{def}}{=} \frac{D_{A2}}{D_{A1}}.
$$

and thus the reaction-diffusion matrix we need to test for the necessary condition $\boldsymbol{\mathcal{N}}$ takes the form

$$\widetilde{\mathbf{M}}(q^2) = \begin{pmatrix} -1-q^2 & 0 & H_{S_{A1}\to A1} & H_{S_{A2}\to A1} \\ 0 & -\delta_{A2/A1} - D_{A2/A1}\cdot q^2 & \delta_{A2/A1}H_{S_{A1}\to A2} & 0 \\ \delta_{S_{A1}/A1} & 0 & -\delta_{S_{A1}/A1} & 0 \\ 0 & \delta_{S_{A2}/A1} & 0 & -\delta_{S_{A2}/A1} \end{pmatrix}.$$

##### 3.4.3 Deriving the necessary condition $\boldsymbol{\mathcal{N}}$ for pattern formation

We compute the determinant of $\widetilde{\mathbf{M}}$:

$$(-1)^n det(\widetilde{\mathbf{M}}(q^2)) = D_{A2/A1}\cdot\delta_{S_{A1}/A1}\cdot\delta_{S_{A2}/A1}\cdot q^4 + \left(\delta_{S_{A1}/A1}\delta_{S_{A2}/A1}\left(D_{A2/A1}(1-H_{S_{A1}\to A1}) + \delta_{A2/A1}\right)\right)\cdot q^2 + \\ + \delta_{S_{A1}/A1}\delta_{S_{A2}/A1}\delta_{A2/A1}\left(1 - H_{S_{A1}\to A1} - H_{S_{A1}\to A2}\cdot H_{S_{A2}\to A1}\right). \tag{36}$$

To satisfy the necessary condition $\boldsymbol{\mathcal{N}}$, the coefficient corresponding to $q^2$ in expression 36 must be negative, i.e.,

$$H_{S_{A1}\to A1} > 1. \tag{37}$$

We also require $a_0 > 0$, implying $1 - H_{S_{A1}\to A1} > H_{S_{A1}\to A2}\cdot H_{S_{A2}\to A1}$. Together, these inequalities imply that $H_{S_{A1}\to A2}\cdot H_{S_{A2}\to A1}$ must be negative and therefore exactly one of the pair $H_{S_{A1}\to A2}, H_{S_{A2}\to A1}$ must be negative. We visualize this via light blue lines in the figures of the main text.

This shows that the model can satisfy the necessary condition $\boldsymbol{\mathcal{N}}$ only if one of the signalling pathways activates the expression of its own signalling (we have chosen this to be the pathway 1), and exactly one pathway inhibits the other (either pathway 1 inhibits pathway 2 or vice versa).

### 4 Parameter values and simulation scripts

All code used to generate the results in this article is available at: https://zenodo.org/records/15495853. It is organised into folders based on the figure numbers.

For easy-to-use, generalized simulation scripts, we instead recommend our open source Julia package: https://github.com/hiscocklab/ReactionDiffusion.jl.

When simulating concrete PDE systems, we approximated the (typically sigmoidal) intracellular response function by a Hill function

$$F([X]) = k \cdot \frac{[X]^n}{K^n + [X]^n},$$

parameterized by: the maximal transcription rate, $k$; the concentration of $[X]$ that yields half-maximal $F$, $K$; and the Hill coefficient $n$. We let all these parameters vary when performing parameter screens. We also varied other parameters such as degradation rates $\delta$, diffusion rates $D$ or association constants $k_+$.
As the initial conditions, we chose the (numerically computed) steady states, perturbed with random noise of order $10^{-2}$. Throughout, we used zero-flux (von Neumann) boundary conditions.

All parameters used in our analysis can be found in the simulation scripts:
https://zenodo.org/records/15495853
We briefly outline the rationale for our parameter choices here:

- Figure 3: The extracellular parameters were chosen to vary consistently across all models. For example:

$$0.1 \leq \delta_{I/A} \leq 10,$$
$$1 \leq k_{I/A} \leq 10,$$
$$0 \leq n_1 \leq 3,$$
$$0 \leq n_2 \leq 3,$$
$$1 \leq k_{+/A} = k_+/\delta_A \leq 100,$$
$$0.1 \leq K_{I/A} \leq 2,$$
$$0.03 \leq D_{I/A} \leq 30.$$

  Parameters specific to each model were chosen in biologically reasonable ranges and so that each model has distinct intracellular signalling (no two models are equivalent under the Schur complement). For the model of GDF5/NOGGIN signalling, we take parameter values from published work [22], adapting the model published in [23]. Strictly speaking, these models are for SMAD2 signalling. Although the quantitative *in vivo* dynamics of SMAD1 may be different from SMAD2, the same intracellular principles are known to apply to both [24]. Therefore, here we assume the same biophysical parameters to demonstrate PDE systems of this level of complexity may be rigorously simulated by *ReactionDiffusion.jl*.

- Figure 4: Here, parameter ranges were selected in an attempt to fully explore the space of $H_{S\to A}$ and $H_{S\to I}$ constrained by the necessary condition $\boldsymbol{\mathcal{N}}$, therefore allowing for wider parameter variations. For example,

$$0.1 \leq \delta_{I/A} \leq 10,$$
$$0.01 \leq \delta_{S/A} \leq 10,$$
$$0.01 \leq D_{I/A} \leq 100.$$

- Figure 5: Here, the parameters shared among the four models $(\delta_{I/A}, k_{I/A}, K_{I/A}, D_{I/A})$ vary consistently between 0.1 and 10, other ranges were chosen to remain biologically plausible , although they are not based on specific experimental data.

#### 4.1 Robustness measures

In this paper we measured two types of robustness:

##### 4.1.1 Fraction of total parameter sets that break symmetry

A well-studied measure of robustness corresponds to the fraction of parameter sets capable of forming Turing patterns within the overall parameter space [25], [5]. This measure depends on the choice of parameter space and thus varies between analyses.
Fixing the value of a concrete free parameter, we calculated the fraction of parameter sets (measured in %) that are predicted to break symmetry. Specifically, for any concrete parameter value $x$ of the free parameter $X$, we compute

$$\frac{\#\text{ parameter sets with X=x that break symmetry}}{\#\text{ all parameter sets with X=x}} \cdot 100.$$

We plotted the results as heatmaps (Figure S5 right). Each row corresponds to fixing a different parameter, with all other parameters allowed to vary across a wide range.

##### 4.1.2 Robustness to individual parameter perturbations

As an alternative robustness metric, we taken a given parameter set $P$ that undergoes a Turing instability and then perturb, one-at-a-time, each parameter in the model. This then determines how much each parameter can be individually varied without compromising the ability of the model to form patterns.
For detailed algorithm see the function

```
function plot_robustness_barcode(turing_params, params, model, index)
```

in the Julia code for Figure 5 at: https://zenodo.org/records/15495853. In all studied models, we found parameter sets with high robustness to these parameter perturbations, for concrete details see Figure S5.

### 5 Appendix 1: Deriving analytical expressions for Turing instabilities in several standard Turing models

In general, it is often infeasible to analytically derive necessary and sufficient conditions for an arbitrary reaction-diffusion system to undergo a Turing instability. Here, we discuss four well-studied systems for which we do have analytical expressions for these necessary and sufficient conditions, and compare them to the numerical predictions obtained by the *ReactionDiffusion.jl* pipeline.

Recall that we say that a system is pattern-forming if there exist two positive real numbers $q_0, q_1$ such that the system is stable (that is, all eigenvalues have negative real parts) in the interval $[0, q_0) \cup (q_1, \infty)$ and has exactly one positive real eigenvalue in the interval $(q_0, q_1)$. Hence, to determine whether a system is pattern-forming, we would need to derive analytical expressions of each eigenvalue and analyse its sign-change properties. In general, this is not possible: first, expressing eigenvalues in terms of the system parameters requires an analytical solution of the steady states, which is generally not possible. Second, the high number of parameters and mathematically complex expressions for each eigenvalue often do not allow for any useful sign-analysis of the eigenvalues.
However, there exist exceptions in which the derivation of concrete analytical formulae for each eigenvalue leads to reasonably simple and understandable conditions for a system to be pattern-forming, i.e., to satisfy the necessary condition $\boldsymbol{\mathcal{N}}$ as well as sufficient conditions. In this section, we will discuss four such systems.

#### 5.1 A general criterion for Turing instability in a two component reaction-diffusion system

Consider a general two-component reaction-diffusion system involving two components, $u$ and $v$ whose reaction is captured by the functions $f(u, v)$, $g(u, v)$ and diffusion constants are given by $D_u$, $D_v$ respectively:

$$\begin{aligned}\frac{\partial u}{\partial t} &= D_u \nabla^2 u + f(u, v) \\ \frac{\partial v}{\partial t} &= D_v \nabla^2 v + g(u, v)\end{aligned} \tag{38}$$

The, setting $D = D_v/D_u$, the reaction-diffusion matrix becomes

$$\mathbf{M} = \begin{pmatrix} f_u - q^2 & f_v \\ g_u & g_v - Dq^2 \end{pmatrix}. \tag{39}$$

Assuming that we can find the analytical solution for the steady state, we are able to directly compute eigenvalues, which are the solutions of

$$\lambda^2 - \lambda(\text{trace}(\mathbf{M})) + \det(\mathbf{M}) = 0 \tag{40}$$

and so may be expressed as

$$\lambda_{\pm}(q) = \frac{1}{2}\left[\text{trace}(\mathbf{M}) \pm \sqrt{([\text{trace}(\mathbf{M})]^2 - 4\det[\mathbf{M}])}\right]. \tag{41}$$

Standard analysis (for details, see for example [26]) reveals that the following four conditions are necessary and sufficient for the $2 \times 2$ reaction-diffusion system to be pattern-forming:

$$\begin{aligned} f_u + g_v &< 0 \\ f_u g_v - f_v g_u &> 0 \\ D f_u + g_v &> 0 \\ \frac{(D f_u + g_v)^2}{4D} + f_v g_u - f_u g_v &> 0 \end{aligned} \tag{42}$$

##### 5.1.1 Gierer-Meinhardt system of equations:

In the Gierer-Meinhardt system of reaction-diffusion equations [19] [9], we have

$$\begin{aligned} f(u,v) &= \frac{u^2}{v} - u + \sigma \\ g(u,v) &= k(u^2 - v), \end{aligned} \tag{43}$$

where $k > 0$.
Using the conditions in (42) we obtain the conditions for this system to be pattern-forming:

$$\begin{aligned} \frac{2\bar{u}}{\bar{v}} - k - 1 &< 0 \\ \frac{k\left(2\bar{u}^3 - 2\bar{u}\bar{v} + \bar{v}^2\right)}{\bar{v}^2} &> 0 \\ \frac{2\bar{u} - \bar{v}}{d\bar{v}} - k &> 0 \\ \left(\frac{2\bar{u} - \bar{v}}{d\bar{v}} - k\right)^2 - \frac{4k\left(2\bar{u}^3 - 2\bar{u}\bar{v} + \bar{v}^2\right)}{d\bar{v}^2} &> 0 \end{aligned} \tag{44}$$

where $d \stackrel{\text{def}}{=\!=} 1/D$ by convention, and $\bar{u}, \bar{v}$ are the steady states of the system when the diffusion is equal to zero. It is easy to see that the steady states may be computed as

$$\bar{u} = 1 + \sigma, \qquad \bar{v} = \bar{u}^2 \tag{45}$$

Thus the conditions that need to be satisfied for the system to be pattern-forming are given by

$$\frac{2}{1+\sigma} - k - 1 < 0 \tag{46a}$$

$$k > 0 \tag{46b}$$

$$\frac{(1-\sigma)}{d(1+\sigma)} - k > 0 \tag{46c}$$

$$\left(\frac{1-\sigma}{d(1+\sigma)} - k\right)^2 - \frac{4k}{d} > 0 \tag{46d}$$

This can be further simplified: for any given $d$, we can find the $k - \sigma$ subspace which satisfies necessary and sufficient conditions for pattern formation:
Notice that the condition in (46)(a) gives that $k > \frac{1-\sigma}{1+\sigma}$. Thus, (46)(a) and (46)(c) could be satisfied simultaneously only if $d < 1$ (that means $D_v > D_u$). Therefore, the lower bound for $k$ is given by the equation

$$k_\ell = \frac{1-\sigma}{1+\sigma} \tag{47}$$

To find the upper bound, we use (46)(d) and solve the quadratic equation, $\left(\frac{1-\sigma}{d(1+\sigma)} - k\right)^2 - \frac{4k}{d} = 0$ and obtain the two possible roots

$$k_\pm = \frac{2}{d} + \frac{1-\sigma}{d(1+\sigma)} \pm \frac{2}{d}\sqrt{\frac{2}{1+\sigma}} \tag{48}$$

The positive root is not acceptable here as that would violate (46)(c). Thus, the upper bound for $k$ is given by

$$k_{\mathcal{U}} = \frac{2}{d} + \frac{1-\sigma}{d(1+\sigma)} - \frac{2}{d}\sqrt{\frac{2}{1+\sigma}} \tag{49}$$

These upper and lower bounds, $k_\ell$ and $k_{\mathcal{U}}$, for $D_v = 1$ and $D_u = 0.1$, are plotted in the Supplementary Figure S1-C. The analytically defined pattern-forming parameter space closely matches the numerical predictions obtained by *ReactionDiffusion.jl*.

###### 5.1.2 The Schnakenberg model:

The Schnakenberg system [27] [7] is defined as

$$\begin{aligned}\frac{\partial u}{\partial t} &= \nabla^2 u + \left(a - u + u^2 v\right)\\ \frac{\partial v}{\partial t} &= D\nabla^2 v + \left(b - u^2 v\right)\end{aligned} \tag{50}$$

For the Schnakenberg system of reaction-diffusion equations, the steady state solutions in absence of diffusion is given by

$$\bar{u} = a + b, \qquad \bar{v} = \frac{b}{\bar{u}^2} = \frac{\bar{u} - a}{\bar{u}^2} \tag{51}$$

Then at $(u, v) = (\bar{u}, \bar{v})$, we have

$$f_u = (2\bar{u}\bar{v} - 1),\ f_v = \bar{u}^2,\ g_u = -2\bar{u}\bar{v},\ g_v = -\bar{u}^2 \tag{52}$$

The first condition from (42) translates to

$$f_u + g_v < 0 \implies 2\bar{u}\bar{v} - 1 - \bar{u}^2 < 0 \implies 1 - \frac{2a}{\bar{u}} - \bar{u}^2 < 0 \tag{53}$$

In order to calculate the lower bound $a_\ell$ from the inequality in (53), we express $\bar{u}, \bar{v}$ in terms of $a$ and $b$ and solve the equation

$$1 - \frac{2a}{(a+b)} - (a+b)^2 = 0. \tag{54}$$

This is a third-degree polynomial in $a$, with exactly one real root. This real root defines the lower bound:

$$a_\ell = -b + \frac{1}{3^{1/3}\left(-9b + \sqrt{3 + 81b^2}\right)^{1/3}} - \frac{\left(-9b + \sqrt{3 + 81b^2}\right)^{1/3}}{3^{2/3}} \tag{55}$$

The second condition from (42) gives

$$f_u g_v - f_v g_u > 0 \implies \bar{u}^2 > 0 \tag{56}$$

which is trivially satisfied. The third condition from (42) translates to

$$Df_u + g_v > 0 \implies D(2\bar{u}\bar{v} - 1) - \bar{u}^2 > 0 \tag{57}$$

Notice that the inequalities in (53) and (57) could be simultaneously satisfied only if $D > 1$. Now, in order to calculate the upper bound $a_{\mathcal{U}}$ we simplify the fourth condition in (42) to obtain

$$\begin{aligned}(Df_u + g_v)^2 - 4D(f_u g_v - f_v g_u) > 0 &\implies \bar{u}^2(D - \bar{u}^2)^2 - 4\bar{u}(D - \bar{u}^2)aD + 4a^2D^2 - 4D\bar{u}^4 > 0\\ &\implies c_1 a^2 + c_2 a + c_3 > 0\end{aligned} \tag{58}$$

where,

$$c_1 = 4D^2, \quad c_2 = -4D\bar{u}(D - \bar{u}^2), \quad c_3 = \bar{u}^2(D - \bar{u}^2)^2 - 4D\bar{u}^4. \tag{59}$$

The equation $c_1 a^2 + c_2 a + c_3 = 0$ has two roots given by

$$a_\pm = \frac{-c_2 + \sqrt{c_2^2 - 4c_1c_3}}{2c_1} = \frac{\bar{u}}{2}\left[1 - \frac{\bar{u}^2}{D} \pm \frac{2\bar{u}}{\sqrt{D}}\right] \tag{60}$$

Since $a_+$ would violate (57), we obtain

$$a_{\mathcal{U}} = a_- = \frac{\bar{u}}{2}\left[1 - \frac{\bar{u}^2}{D} - \frac{2\bar{u}}{\sqrt{D}}\right] \tag{61}$$

Using the fact that $\bar{u} = a + b$, we solve the equation

$$a = \frac{(a+b)}{2}\left[1 - \frac{(a+b)^2}{D} - \frac{2(a+b)}{\sqrt{D}}\right] \tag{62}$$

This equation has three roots, one real and two complex. The only real root gives the value of $a_{\mathcal{U}}$ :

$$a_{\mathcal{U}} = \frac{1}{3}\left[-3b - 2\sqrt{D} + \frac{D}{\left(27bD + D^{3/2} + \sqrt{729b^2D^2 + 54bD^{5/2}}\right)^{1/3}} + \left(27bD + D^{3/2} + \sqrt{729b^2D^2 + 54bD^{5/2}}\right)^{1/3}\right]. \tag{63}$$

These upper and lower bounds, $a_\ell$ and $a_{\mathcal{U}}$, for $D_v = 50$ and $D_u = 1$, are plotted in the Supplementary Figure S1-D. The analytically defined pattern-forming parameter space closely matches the numerical predictions obtained by *ReactionDiffusion.jl*.

###### 5.1.3 The CIMA model

In the CIMA model [28] [8], we have

$$\begin{aligned}\frac{\partial u}{\partial t} &= D_u\nabla^2 u + a - u - \frac{4uv}{1+u^2}\\ \frac{\partial v}{\partial t} &= D_v\nabla^2 v + \sigma b\left(u - \frac{uv}{1+u^2}\right),\end{aligned} \tag{64}$$

where $\sigma, b, \alpha > 0$. For the CIMA system of reaction-diffusion equations, the steady state solutions are given by

$$\bar{u} = \frac{a}{5} \stackrel{\text{def}}{=\!=} \alpha, \qquad \bar{v} = 1 + \frac{a^2}{25} = 1 + \alpha^2. \tag{65}$$

Then, at $(u, v) = (\bar{u}, \bar{v})$, we have

$$f_u = \frac{\partial f}{\partial u} = \frac{3\alpha^2 - 5}{1+\alpha^2},\ f_v = \frac{\partial f}{\partial v} = -\frac{4\alpha}{1+\alpha^2},\ g_u = \frac{\partial g}{\partial u} = \sigma b\left(\frac{\alpha^2}{1+\alpha^2}\right),\ g_v = \frac{\partial g}{\partial v} = -\sigma b\frac{\alpha}{1+\alpha^2}. \tag{66}$$

Following (42), the CIMA system is pattern-forming iff the following conditions are satisfied,

$$3\alpha^2 - 5 < \sigma b\alpha \tag{67a}$$

$$\sigma b\alpha(1+\alpha^2) > 0 \tag{67b}$$

$$D(3\alpha^2 - 5) > \sigma b\alpha \tag{67c}$$

$$D^2(3\alpha^2 - 5)^2 - 26\alpha^3 D\sigma b - 10D\sigma b\alpha + \sigma^2 b^2\alpha^2 > 0 \tag{67d}$$

Notice that (67a) and (67c) could be simultaneously satisfied only if $D > 1$.
We fix $\sigma = 1$ and then find the boundary limits of the pattern-forming space. The lower bound is obtained from (67a) and is given by

$$b_\ell = \frac{3a}{5} - \frac{25}{a}. \tag{68}$$

In order to find the upper-bound on the values of $b$, we solve the equation

$$D^2\left(3\alpha^2 - 5\right)^2 - 26\alpha^3 Db - 10Db\alpha + b^2\alpha^2 = 0,$$

and obtain the two roots of $b$:

$$b_\pm = \frac{D}{5a}\left[125 + 13a^2 \pm 4a\sqrt{10(a^2+25)}\right]. \tag{69}$$

The root $b_+$ is not acceptable as it violates (67c). Therefore the upper bound is given by

$$b_{\mathcal{U}} = \frac{D}{5a}\left[125 + 13a^2 - 4a\sqrt{10(a^2+25)}\right]. \tag{70}$$

These upper and lower bounds, $b_\ell$ and $b_{\mathcal{U}}$, for $D_v = 15$ and $D_u = 1$, are plotted in the Supplementary Figure S1-E. The analytically defined pattern-forming parameter space closely matches the numerical predictions obtained by *ReactionDiffusion.jl*.

##### 5.2 Generalized 4-component systems

Some analytical results for necessary and sufficient conditions for Turing instability in 4-component reaction-diffusion systems have been obtained in [10], where the authors studied the effect of coupling between two two-component reaction diffusion systems. A generalized system can be written as

$$\frac{\partial u}{\partial t} = \gamma f(u, v, \tilde{u}, \tilde{v}) + \nabla^2 u \tag{71a}$$

$$\frac{\partial v}{\partial t} = \gamma g(u, v, \tilde{u}, \tilde{v}) + D\nabla^2 v \tag{71b}$$

$$\frac{\partial \tilde{u}}{\partial t} = \gamma f(\tilde{u}, \tilde{v}, u, v) + \nabla^2 \tilde{u} \tag{71c}$$

$$\frac{\partial \tilde{v}}{\partial t} = \gamma g(\tilde{u}, \tilde{v}, u, v) + D\nabla^2 \tilde{v} \tag{71d}$$

where $\gamma > 0$ and $D > 0$ is the diffusion parameter. Then the reaction-diffusion matrix takes the form

$$\mathbf{M} = \begin{bmatrix} f_u - q^2 & f_v & f_{\tilde{u}} & f_{\tilde{v}} \\ g_u & g_v - Dq^2 & g_{\tilde{u}} & g_{\tilde{v}} \\ f_{\tilde{u}} & f_{\tilde{v}} & f_u - q^2 & f_v \\ g_{\tilde{u}} & g_{\tilde{v}} & g_u & g_v - Dq^2 \end{bmatrix} \tag{72}$$

Let $(\bar{u}, \bar{v}, \tilde{\bar{u}}, \tilde{\bar{v}})$ be the steady state in the absence of diffusion. Performing the analysis analogous to the $2 \times 2$ system (the characteristic polynomial may be factorized as a product of two second-order polynomials), we find the following conditions for a system to be pattern-forming: for Turing instability to occur one of the following sets of conditions needs to be satisfied

$$f_u + g_v < \pm(f_{\tilde{u}} + g_{\tilde{v}}), \tag{73a}$$

$$f_u g_v - f_v g_u + f_{\tilde{u}} g_{\tilde{v}} - f_{\tilde{v}} g_{\tilde{u}} > \pm(f_u g_{\tilde{v}} - f_{\tilde{v}} g_u + f_{\tilde{u}} g_v - f_v g_{\tilde{u}}), \tag{73b}$$

$$D f_u + g_v + D f_{\tilde{u}} + g_{\tilde{v}} > 0, \tag{73c}$$

$$(D f_u + g_v + D f_{\tilde{u}} + g_{\tilde{v}})^2 - 4D(f_u g_v - f_v g_u + f_u g_{\tilde{v}} - f_{\tilde{v}} g_u + f_{\tilde{u}} g_v - f_v g_{\tilde{u}} + f_{\tilde{u}} g_{\tilde{v}} - f_{\tilde{v}} g_{\tilde{u}}) > 0; \tag{73d}$$

$$f_u + g_v < \pm(f_{\tilde{u}} + g_{\tilde{v}}), \tag{74a}$$

$$f_u g_v - f_v g_u + f_{\tilde{u}} g_{\tilde{v}} - f_{\tilde{v}} g_{\tilde{u}} > \pm(f_u g_{\tilde{v}} - f_{\tilde{v}} g_u + f_{\tilde{u}} g_v - f_v g_{\tilde{u}}), \tag{74b}$$

$$D f_u + g_v - D f_{\tilde{u}} - g_{\tilde{v}} > 0, \tag{74c}$$

$$(D f_u + g_v - D f_{\tilde{u}} - g_{\tilde{v}})^2 - 4D(f_u g_v - f_v g_u - f_u g_{\tilde{v}} + f_{\tilde{v}} g_u - f_{\tilde{u}} g_v + f_v g_{\tilde{u}} + f_{\tilde{u}} g_{\tilde{v}} - f_{\tilde{v}} g_{\tilde{u}}) > 0, \tag{74d}$$

For detailed derivation see [10].

###### 5.2.1 The C1 model:

The C1 model is a 4-component reaction-diffusion system [10], defined as

$$f = a - u + u^2 v + p\tilde{u}^2 \tilde{v} \tag{75a}$$

$$g = b - u^2 v \tag{75b}$$

$$\tilde{f} = a - \tilde{u} + \tilde{u}^2 \tilde{v} + p u^2 v \tag{75c}$$

$$\tilde{g} = b - \tilde{u}^2 \tilde{v}. \tag{75d}$$

For the C1 model, the steady state solution in the absence of diffusion is given by

$$\begin{aligned}
\bar{u} &= a + b + pb \\
\bar{v} &= \frac{b}{\bar{u}^2} \\
\bar{\bar{u}} &= a + b + pb \\
\bar{\bar{v}} &= \frac{b}{\bar{\bar{u}}^2}
\end{aligned} \tag{76}$$

The lower bound can be obtained by satisfying the conditions in (73a) and (74a) which gives

$$2\bar{u}\bar{v} - 1 - \bar{u}^2 < \pm\,(2p\bar{\bar{u}}\bar{\bar{v}})\,. \tag{77}$$

From (76) it follows that $\bar{u} = \bar{\bar{u}}$ and $\bar{v} = \bar{\bar{v}}$. Thus in (77), the inequality is simplified as

$$2\bar{u}\bar{v} + 2p\bar{u}\bar{v} - 1 - \bar{u}^2 < 0 \tag{78}$$

Solving $\bar{u}^2 - 2(1+p)\bar{u}\bar{v} + 1 = 0$, with the values of $\bar{u}$ and $\bar{v}$ given in (76), the lower bound is given by

$$a_\ell = \frac{h(b)}{3^{2/3}\cdot 10^{1/3}} - \frac{11b}{10} - \frac{10^{1/3}}{3^{1/3}h(b)} \tag{79}$$

where

$$h(b) \stackrel{\text{def}}{=\!=} (99b + 3^{1/2}(3267b^2 + 100)^{1/2})^{1/3}. \tag{80}$$

Choosing $p = 0.1$, the upper bound can be obtained by satisfying the conditions in (73d) which gives

$$(Df_u + g_v + Df_{\bar{\bar{u}}} + g_{\bar{\bar{v}}})^2 - 4D(\bar{u}^2) > 0 \tag{81}$$

This is lucky, because we can simply use formula for $A^2 - B^2$ to split sixth-degree polynomial into product of two third-degree polynomials:

$$(Df_u + g_v + Df_{\bar{\bar{u}}} + g_{\bar{\bar{v}}} - 4\sqrt{D}\bar{u}) \cdot (Df_u + g_v + Df_{\bar{\bar{u}}} + g_{\bar{\bar{v}}} + 4\sqrt{D}\bar{u}) > 0 \tag{82}$$

Solving $Df_u + g_v + Df_{\bar{\bar{u}}} + g_{\bar{\bar{v}}} - 4\sqrt{D}\bar{u} = 0$, with the values of $\bar{u}$ and $\bar{v}$ given in (76), the upper bound is given by

$$a_u = \frac{10^{2/3}h_2(b,D)}{30} - \frac{11b}{10} - \frac{2D^{1/2}}{3} + \frac{10^{1/3}tD}{3h_2(b,D)} \tag{83}$$

where

$$h_2(b,D) \stackrel{\text{def}}{=\!=} (297bD + 10D^{3/2} + 3\cdot 33^{1/2}\cdot(bD^2(297b + 20D^{1/2}))^{1/2})^{1/3}. \tag{84}$$

These upper and lower bounds, $a_\ell$ and $a_u$, for $D_v = 50$ and $D_u = 1$, are plotted in the Supplementary Figure S1-F. The analytically defined pattern-forming parameter space closely matches the numerical predictions obtained by *ReactionDiffusion.jl*.
